# Social cognitive regions of human association cortex are selectively connected to the amygdala

**DOI:** 10.1101/2023.12.06.570477

**Authors:** Donnisa Edmonds, Joseph J. Salvo, Nathan Anderson, Maya Lakshman, Qiaohan Yang, Kendrick Kay, Christina Zelano, Rodrigo M. Braga

## Abstract

Reasoning about someone’s thoughts and intentions – i.e., forming a theory of mind – is an important aspect of social cognition that relies on association areas of the brain that have expanded disproportionately in the human lineage. We recently showed that these association zones comprise parallel distributed networks that, despite occupying adjacent and interdigitated regions, serve dissociable functions. One network is selectively recruited by theory of mind processes. What circuit properties differentiate these parallel networks? Here, we show that social cognitive association areas are intrinsically and selectively connected to regions of the anterior medial temporal lobe that are implicated in emotional learning and social behaviors, including the amygdala at or near the basolateral complex and medial nucleus. The results suggest that social cognitive functions emerge through coordinated activity between amygdala circuits and a distributed association network, and indicate the medial nucleus may play an important role in social cognition in humans.

## Introduction

The ability to reason about another person’s intentions and beliefs – i.e., to form a ‘theory of mind’ (ToM) – is an important aspect of social cognition that assists the navigation of social groups.^1^ In humans, tasks targeting ToM activate a set of association regions of the brain that are late to mature and are disproportionately expanded in the hominin lineage,^2–5^ supporting that the primate brain may have expanded following evolutionary pressures associated with living in complex social groups.^6^ Additionally, evidence supports that evolutionarily ancient structures, such as the amygdala and related anterior medial temporal lobe (MTL) circuitry, are central to social behaviors. It is unclear how these lines of evidence intersect. Here, we show that human association regions that are active during ToM are intrinsically and selectively connected to the amygdala, including nuclei that are extensively implicated in social behaviors in rodents.^7–10^

Brain regions involved in ToM can be examined in humans by studying task-related changes in the blood-oxygenation-level-dependent (BOLD) signal^11,12^ during ‘false belief’ and ‘emotional pain’ tasks. In the false belief task,^13,14^ participants answer questions from the perspective of someone who holds a mistaken belief (e.g., Sally believes a cherry cake tastes of strawberries because it was mislabeled). In the ‘emotional pain’ task, participants rate the pain a protagonist may feel following an emotionally painful event (e.g., the loss of a family pet).^15–17^ Both tasks require thinking about someone else’s thoughts and activate a network of cortical regions that includes the temporoparietal junction, the ventromedial and dorsomedial prefrontal cortex, the lateral temporal cortex, and posteromedial cortex.^16,18–20^ However, early observations noted that these same regions were recruited during other tasks,^21–24^ including those targeting autobiographical memory,^25^ self-oriented thinking,^23,24,26–28^ and ‘episodic projection’ (EP; i.e., thinking about the past or future).^29–31^ The same set of association regions was also conceptualized as the canonical ‘default network’ (DN),^21,32,33^ which is definable from resting-state correlations of the BOLD signal (i.e., functional connectivity or FC),^34,35^ exhibits connectivity to the MTL,^21,35–37^ and shows increased activity during resting periods between active tasks.^38^ This overlap between ToM, the DN, and other memory-related processes led to the idea that the DN plays a domain-general role in introspection and mind-wandering, which tends to include thoughts about others^29,39–47^ (see twelfth figure in Buckner et al.^21^ and first figure in Mars et al.^22^).

An alternative view is that the canonical DN appears to be domain-generalized because it is a coarse (i.e., blurry) estimate of finer-grained, domain-specialized networks. For instance, it was noted that even in group-averaged estimates, which tend to result in blurring over individual differences in anatomy,^48–51^ there are hints of substructure within the canonical DN;^41,52–59^ ToM tasks tend to activate a more anterior region of the inferior parietal lobe at or near the temporoparietal junction, while EP tasks recruit more posterior regions.^55,57,58^ Similarly, ToM tasks typically recruit dorsal posteromedial regions at or near the posterior cingulate cortex, while EP tasks typically recruit more ventral regions at or near the retrosplenial cortex. These distinctions presaged findings from finer-grained, individual-focused analyses. Within individuals, separable posteromedial regions were found to be active when participants are asked to think about relationships between people or places.^20,60–63^ This suggests a separation between social and episodic processes within the DN, and that EP-related activity is likely related to spatial mnemonic processes or mental ‘scene construction.’^47,64,65^ In these examples, social cognition was again located to more dorsal posteromedial regions, whereas EP-related activity was located in more ventral regions, matching the distinctions in group-averaged data. These findings indicate that the convergence of functions on the canonical DN may have been due to blurring across distinct, adjacent regions which separately support social and episodic functions.^24^

Recently, individual-level estimates of brain networks have been achieved by performing FC on extensively sampled individuals.^66–69^ We showed that at least two distinct networks can be identified within the canonical DN, and that the networks display adjacent, interdigitated regions in multiple association zones, leading to the appearance of parallel distributed networks.^66,70^ Pertinently, the two networks, termed “DN-A” and “DN-B”, recapitulated the aforementioned dorsal-ventral distinction in the posteromedial cortex, as well as the anterior-posterior distinction in the inferior parietal lobe, between ToM- and EP-related activity. Confirming this dissociation, the FC estimates of DN-B closely overlapped with activity evoked by ToM tasks within the same individuals, while the adjacent network, DN-A, instead showed activity during an EP task that involved mental scene construction.^20^ The correspondence between the FC and task-evoked responses was evident throughout the brain, rather than just within the posteromedial and inferior parietal components, suggesting network-level specialization of the parallel networks within the canonical DN.^71^

How do these two adjacent, interdigitated, networks support dissociable functions? And how do the circuit properties of DN-B enable social cognitive functions (i.e., ToM)? An answer may lie in how each network relates to more evolutionary ancient non-neocortical structures.^72^ We previously noted that DN-A contains a region in the posterior parahippocampal cortex^66^ and the subiculum,^70^ which suggests a link between the cortical regions of DN-A and the mnemonic and spatial functions of the posterior MTL.^36,47,73–76^ This link provides an explanation for why association regions that constitute DN-A are active during EP; they work with the hippocampal formation to enable mental scene construction.

Although DN-A and DN-B contain adjacent regions in multiple cortical zones, we did not observe an adjacent region of DN-B in the MTL in our prior estimates. However, there are reasons to think such a region may have been missed. First, anterior potions of the temporal lobe suffer from signal dropout in conventional BOLD imaging,^77^ which may have obscured characterization of this region. Smaller voxel sizes, such as those available at higher field strength (e.g., 7T), can reduce signal dropout.^78^ Second, network regions in the anterior MTL, particularly in the amygdala, may be small and fall beneath the resolution limits of conventional 3T fMRI. We previously showed that high-resolution 7T fMRI can reveal small DN-A regions at or near the subiculum that were missed at 3T^70^ (See also Sladky et al.^79^; Maass et al.^80^; Gorka et al.^81^). Third, there are reports of a link between the amygdala and the canonical DN in group-averaged FC data,^81–85^ though these prior efforts did not differentiate between regions supporting social and episodic processes (i.e., DN-A and DN-B). Finally, a recent investigation at 7T provided evidence for a link between the entorhinal cortex and a network resembling DN- B,^86^ supporting a closer link between DN-B and the anterior MTL.

There is also ample evidence to support a connection between the anterior MTL, specifically the amygdala, and socially relevant information processing.^10,87,88^ Lesions of the amygdala lead to loss of social status in primates and rodents,^89^ as well as decreased social affiliation, inappropriate responses to social cues, and both increases and decreases in avoidance.^7,8,90,91^ Amygdala neurons show increased responses when facial expressions and social interactions are viewed,^92–97^ as well as to eye contact from conspecifics.^98^ In humans, lesions of the amygdala lead to difficulties in recognizing emotions in facial expressions,^91,99–105^ and amygdala responses track the perceived emotion rather than visual properties of viewed expressions.^96^ Furthermore, the amygdala has been implicated in psychiatric disorders that are characterized by disrupted social functioning, including autism, as well as schizophrenia, anxiety, and depression.^106–109^ Thus, the evidence suggests that the amygdala is an evolutionarily conserved structure that is crucial for social behaviors and may work in tandem with cortical regions to enable complex social functions such as ToM.^92,105,110^ These findings motivate a closer investigation of the anterior MTL, including the amygdala, and the distributed association networks using high-resolution 7T fMRI.

## Results

### Discovery and replication strategy for high-resolution 7T fMRI data

This study analyzed high-resolution (1.8-mm isotropic) 7T fMRI resting-state data from 8 extensively sampled participants from the Natural Scenes Dataset (NSD).^111^ Two participants (S5 and S8) were excluded during quality control for excessive head motion, leading to a sample of 6 participants (ages 23-27 years) who provided 6-35 resting-state runs each (S1: 35 runs; S2: 6; S3: 16; S4: 12; S6: 19; and S7: 18). For participants providing 12 or more runs passing quality control, data were divided into 2 or 3 datasets for discovery (i.e., free exploration of FC patterns), replication and triplication within individuals. A temporal signal to noise ratio (tSNR) map confirmed that each participant provided high tSNR despite the small voxel size, including in anterior MTL regions critical to our hypothesis (see **Supp. Fig. S1**).

### Replication of parallel distributed networks within the canonical default network

Using the discovery dataset, we followed Braga and Buckner^66^ to define DN-A and DN-B on surface-projected data^112^ using FC from seeds manually selected from the left dorsal prefrontal cortex. Once candidate seeds were selected, network definition was repeated using a data-driven multi-session hierarchical Bayesian model^113^ (MS-HBM; **Fig. 1**) to produce converging estimates of the networks. This provided another replication of the separation between the networks.^20,70,86,115–118^ Surface-based analysis allows easy observation of the cortical mantle, making it convenient for identifying the networks based on anatomical characteristics (see detailed descriptions in Braga and Buckner^66^ and Braga et al.^70^), but does not well-capture the anatomy of the MTL. Therefore, we re-defined DN-A and DN-B in the volume using manually selected seed voxels from left dorsal prefrontal cortex, and projected the volume-defined maps to the surface for comparison with surface-defined estimates to confirm that both approaches were targeting the same networks (**Fig. 1**). Following Braga and Buckner,^70^ we binarized the correlation maps above a threshold of r > 0.3 to compute an overlap map. In each participant, the primary distinctions between networks DN-A and DN-B, such as in the inferior parietal lobe and posteromedial cortex, were present in both surface and volume estimates (**Fig. 1**).

**Figure 1:**
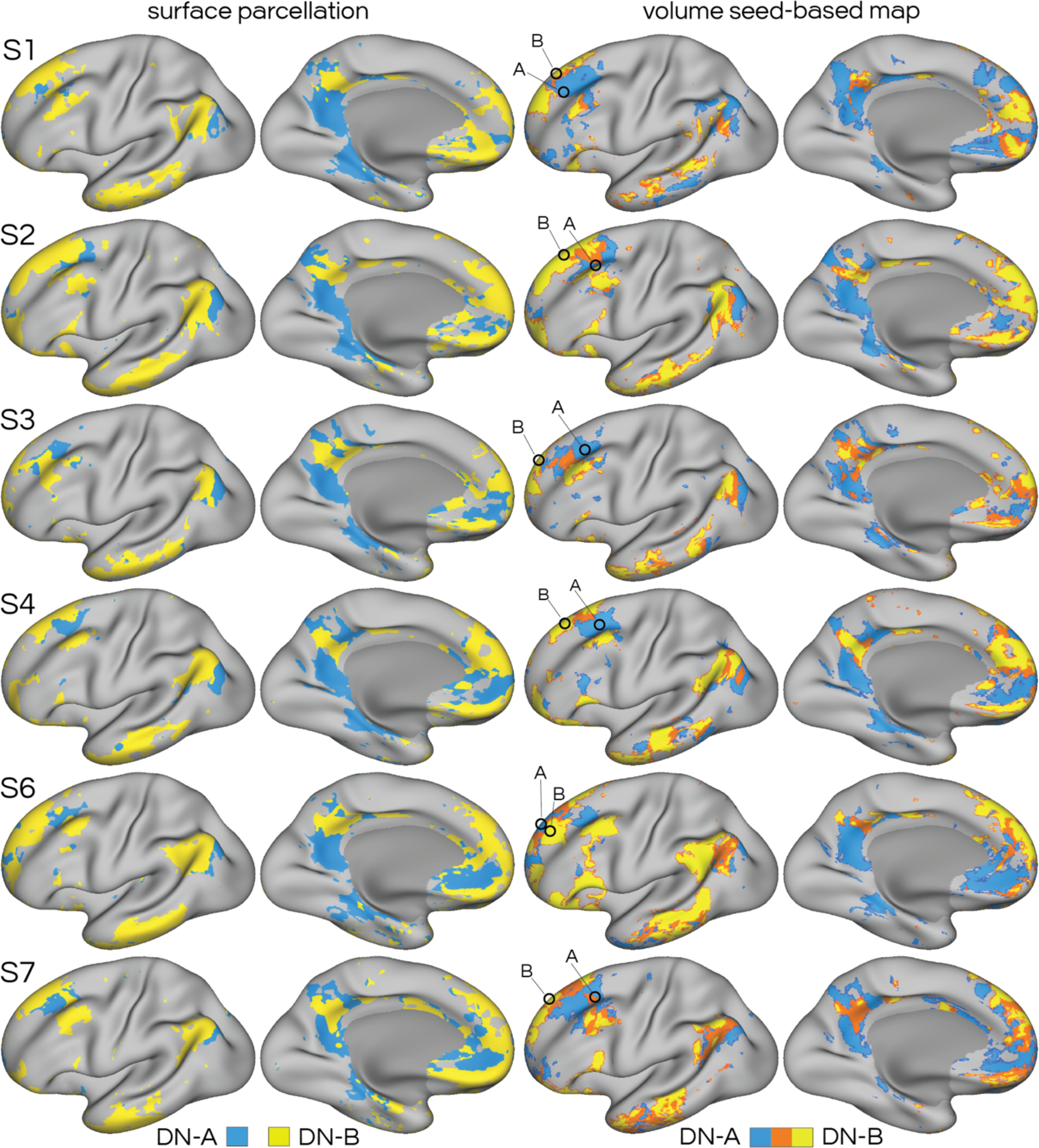
High-resolution functional connectivity (FC) separates default network A (DN-A) and B (DN-B) using both surface- and volume-based approaches. The left column shows the networks defined on surface-projected data for each subject (S1-4, S6, S7) using data-driven clustering^163^ (*k* = 14). The surface visualization allows easier observation of the cortical sheet, but excludes medial temporal structures (see darker gray midline areas). The networks were therefore identified independently in the volume (right column) and projected to the surface for comparison between methods. The volume-defined maps were thresholded at 0.3 and binarized, to display where the estimates of DN-A (blue) and DN-B (yellow) are distinct and overlap (orange). Even in the surface, at high-resolution, DN-B showed evidence of anterior medial temporal regions (see comparison to 3T in Supp. Fig. S2).

### Replication of functional dissociation between DN-A and DN-B

We previously showed that FC estimates of DN-B overlap closely with activity evoked by ToM tasks within individuals.^20^ To build confidence in this link, and support that our estimate of DN-B was delineating association regions related to social cognition, we replicated this result in a separate ‘detailed brain network organization’ (DBNO) dataset collected at Northwestern University. DN-A and DN-B were defined at 3T, using the same procedures as those used in the NSD data, in 8 individuals who also provided ToM (i.e., false-belief and emotional pain) and EP (i.e., mental ‘scene construction’) task data. **Supp. Fig. S2** allows side-by-side comparison of maps of DN-A and DN-B defined in the 7T and 3T subjects, to show that the same networks are being studied in each dataset. In all 8 DBNO subjects, we observed selective overlap between DN-B and ToM-active regions (**Fig. 2**). Conversely, DN-A overlapped with regions active during mental scene construction. This confirms that the FC estimates of DN-B encapsulates regions that are specialized for social cognitive functions and are functionally dissociated from DN-A.

**Figure 2:**
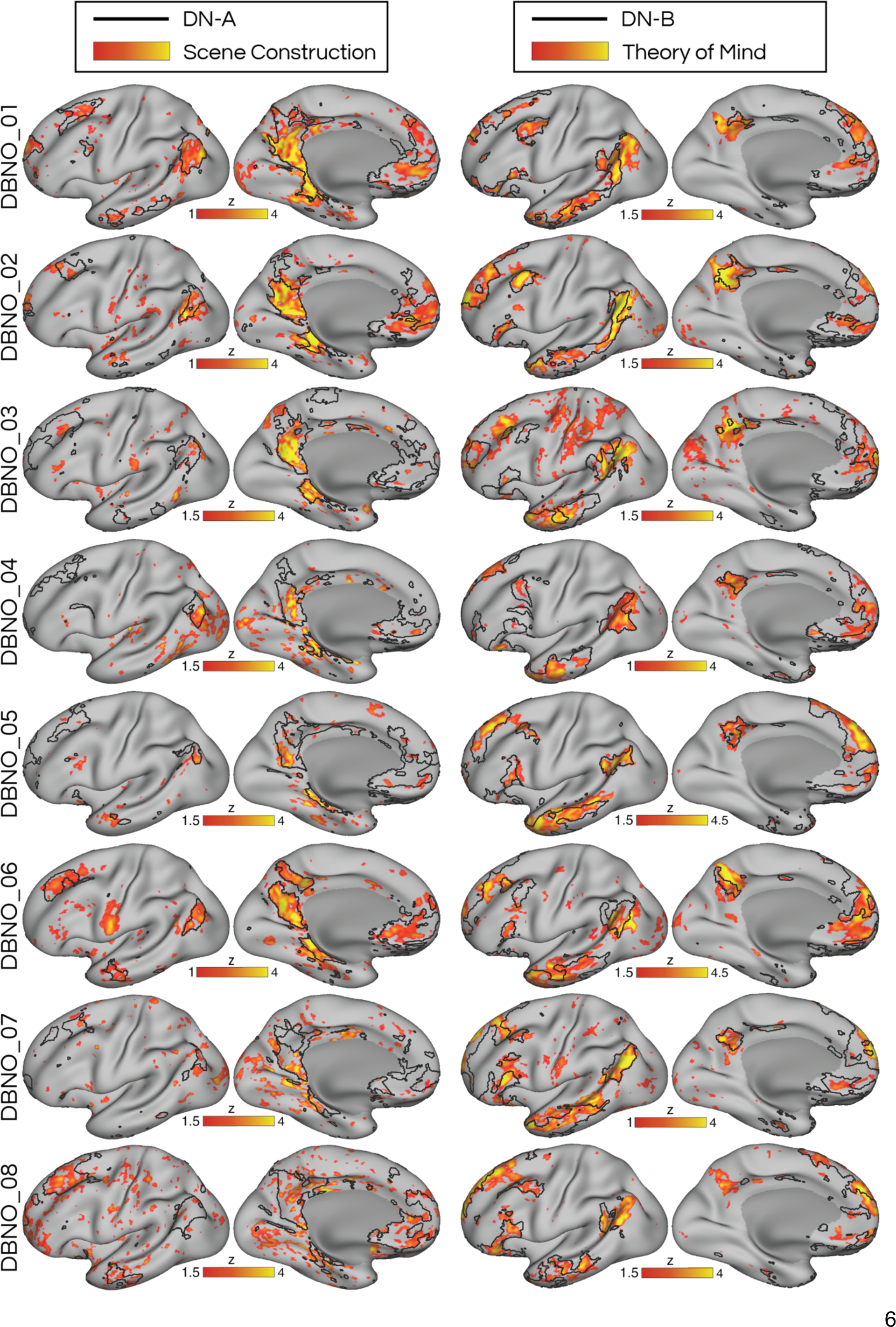
**Functional imaging at 3T in eight additional individuals (DBNO_01–DBNO_08) confirms functional dissociation between DN-A and DN-B**. Black borders show the FC estimates of DN-A (left) and DN-B (right; from **Supp. Fig. S2)**. The same participants provided data during tasks targeting theory of mind (ToM) and mental scene construction. The ToM task contrasts revealed increased activity in multiple regions that overlapped selectively with the boundaries of DN-B. In contrast, the scene construction contrast, which targeted episodic projection (EP), revealed that while participants imagined scenes, increased activity was evident in regions that overlapped with DN-A. The results confirm^20^ that network DN-B is selectively recruited during social cognitive processes, and is functionally dissociated from DN-A.

### Discovery that DN-B contains bilateral regions in the amygdala

The surface-based estimates of DN-B from the 7T fMRI data displayed hints of regions in the anterior MTL not clearly evident at 3T (**Supp. Fig. S2**; and see other 3T estimates in second, ninth and seventeenth figures in DiNicola^20^). To study the anterior MTL more closely, we examined the volume-defined estimates at a correlation threshold of r > 0.2, following Braga and Buckner ^66^ and Braga et al.^70,119^, accounting for the lower tSNR in these regions (**Supp. Fig. S1**). **Fig. 3** shows sagittal views of DN-A and DN-B, along with the overlap map, in participant S2. Confirming our previous findings,^70^ DN-A displayed a prominent posterior MTL region at or near the parahippocampal cortex and a further smaller region at or near the subiculum. We discovered that DN-B displays multiple regions in the anterior MTL: one putatively in the entorhinal cortex,^86^ but intriguingly, other regions were also observed that appeared to be in the amygdala (**Fig. 3**). Regions of DN-A and DN-B were located in mostly non-overlapping locations, even at the lower correlation threshold of 0.2. The putative amygdala regions displayed lower correlation values than the non-amygdala regions in this subject. To confirm these regions were not due to noise, we examined a coronal slice approximately around MNI y = -6 to -9, in each subject (**Fig. 4**). This revealed that the putative amygdala regions of DN-B were bilateral in 4/6 subjects, and likely not spurious. Subject S1 displayed unilateral regions, while S6 did not show a DN-B region that was distinct from DN-A at this location. For these two subjects, we lowered the threshold to 0.15, after which participant S1 showed clearer evidence of bilateral regions (**Fig. 4**), but subject S6 still showed no reliable regions (see also replication and triplication analyses in **Supp. Fig. S6**). Nonetheless, the analyses provided evidence that distributed network DN-B contains regions within the human amygdala that are bilateral and identifiable in multiple individuals.

**Figure 3:**
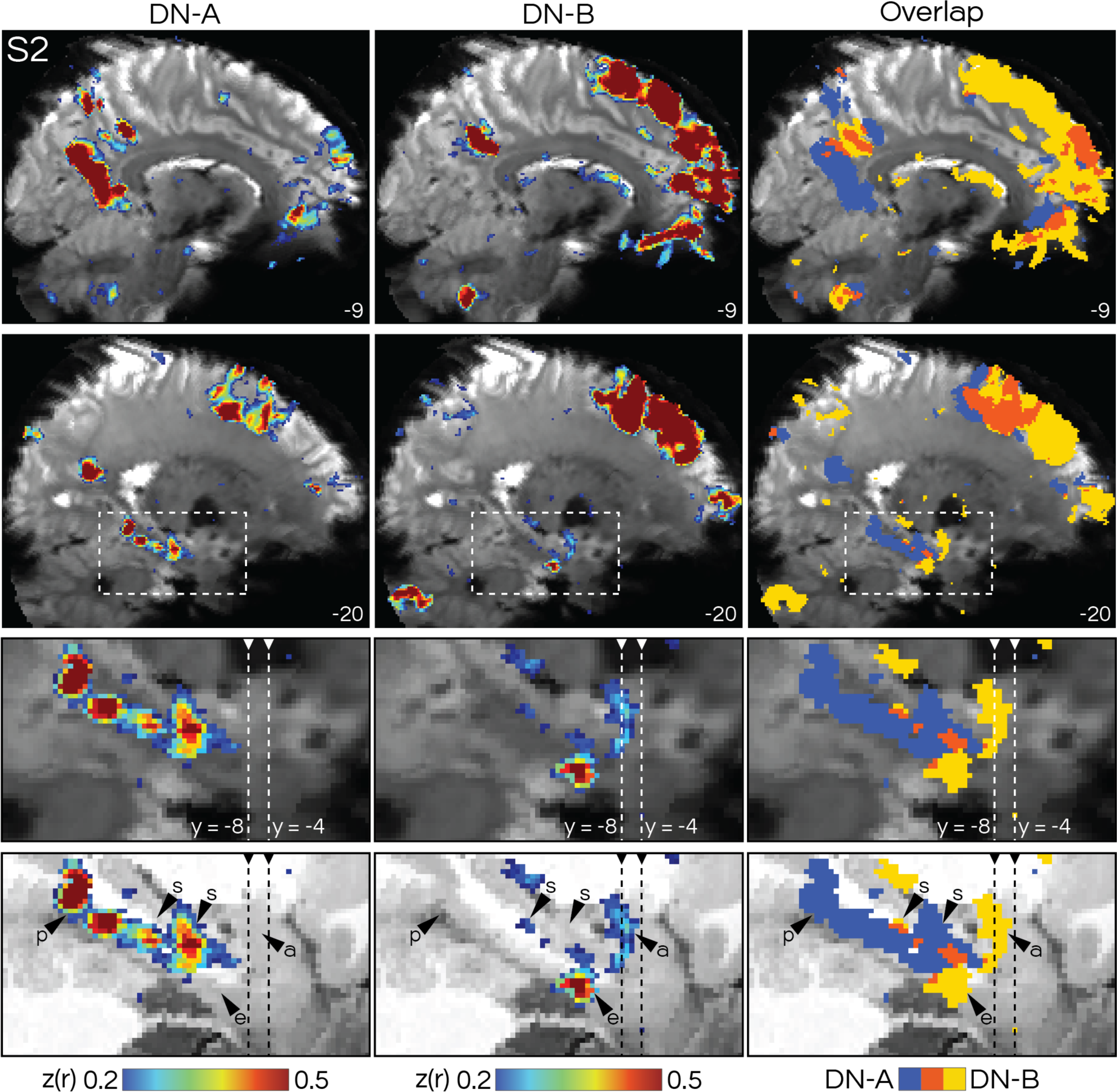
Network DN-B contains distinct regions in the anterior medial temporal lobe (MTL), including amygdala (a), entorhinal cortex (e), and potentially subiculum (s). Sagittal views of the volume-based FC maps of DN-A and DN-B from an example subject (S2) are shown, with the mean blood-oxygenation-level-dependent (BOLD) image as an underlay. The correlation maps were thresholded at 0.2 and binarized to display an overlap map. The top row shows the midline, to display the characteristic posteromedial distinctions between the networks. The second row shows a slice along the MTL, where interdigitated regions of DN-A and DN-B can be seen along the long axis, with DN-B displaying regions in anterior MTL. The dashed boxes indicate the location of the zoom-in shown in the lower two rows. The lowest row shows the T1 image as the underlay for better appreciation of the anatomy. The dashed vertical lines indicate the location of the coronal slices shown in Fig. 4. *z*(*r*), Fisher’s *r*-to-*z* transformed Pearson’s product-moment correlations. Numbers in each panel correspond to MNI slice coordinates. p, parahippocampal cortex.

**Figure 4:**
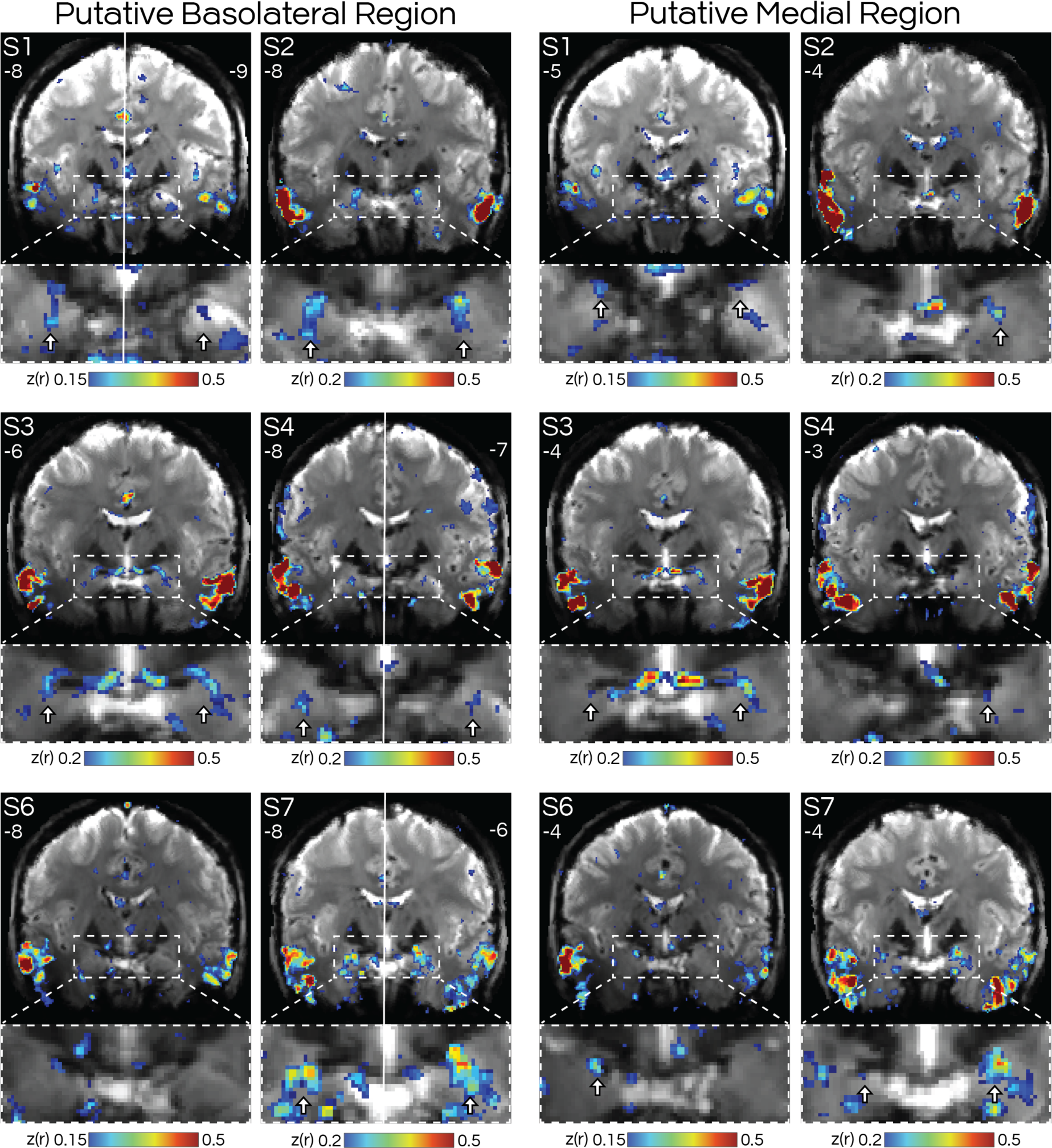
**Basolateral and medial amygdala regions of DN-B are bilateral and replicate across participants**. Volume-based FC maps of DN-B are shown in coronal slices. Numbers refer to MNI coordinate of each slice. On the left are views around y = -6 to -9, where 5 out of 6 subjects (exception: S6) displayed bilateral regions putatively in the basolateral amygdala. On the right are slices around y = -3 to -5, where in all 6 subjects, a distinct set of bilateral (3 out of 6) or unilateral (3 out of 6) regions could be seen putatively near the medial amygdala (Fig. 6). The white solid line in S1, S4 & S7 denotes that left and right hemispheres are from different slices. Arrows denote putative DN- B regions that were distinct from DN-A (see Figs. 5 & 6 for overlap map at these same slices, and replications in **Supp. Figs. S6 & S7**).

### Replication and triplication of amygdala region of DN-B through multiple approaches

We confirmed the results were consistent across multiple analysis approaches. First, in the discovery dataset, a seed voxel was placed directly in the amygdala, targeting the region of DN- B in each subject. The resulting correlation maps were projected to the surface for comparison with the surface-defined networks (**Supp. Fig. S3**). In all subjects, the amygdala seed produced a correlation map that overlapped with the boundaries of DN-B (see especially subjects S2, S4 and S7), despite the low tSNR found around the amygdala. Participants S1 and S3 displayed relatively weak correlations that still overlapped with DN-B. These analyses confirmed that the amygdala is connected to the full distributed network DN-B, and confirmed that this link is selective, with the amygdala-seeded map showing limited overlap with DN-A.

We also replicated our findings using the independent, left-out data from each participant. We re-estimated DN-A and DN-B in the volume in the replication and triplication datasets using dorsal prefrontal seeds (**Supp. Fig. S5**). For each subject, we examined the MNI coordinates where we initially discovered amygdala regions and found amygdala regions of DN-B were replicated in all subjects who displayed a region in the discovery dataset (**Supp Fig. S6**). For one subject, S1, the region replicated in 1 of the 2 left-out datasets. The DN-B regions were located within the same approximate locations, though did vary slightly, with potential reasons discussed in *Limitations and Technical Considerations.* We therefore report with confidence that DN-B contains bilateral amygdala regions that are reproducible within and across participants and generalize across analysis approaches.

### Confirmation that DN-B regions are in the basolateral complex of the amygdala

The amygdala is a complex structure that contains multiple nuclei with distinct roles and connections.^10,96,120^ Detailed knowledge of how these nuclei relate to the extended cortical network involved in social cognition is important. Although it is difficult to study amygdala nuclei in the human brain using neuroimaging, confidence can be built by using individualized estimates of the amygdala anatomy, and focusing on findings that are consistent across individuals. We overlaid the network maps onto automated, individualized segmentations of the amygdala.^112,121^ We also compared our findings to a group-defined amygdala atlas,^120^ which broadly confirmed the results, but with less specificity (not shown). **Fig. 5** and **Supp. Fig. S8** show a zoom-in on the amygdala, displaying how the individualized amygdala segmentation relates to the anatomical T1 and BOLD images from each participant. The demarcations should be interpreted as approximations. For instance, in subject S2 the boundaries of the lateral nucleus appear to overextend the amygdala (**Supp. Fig. S8**). Here, our interpretations are based on a combination of the atlas boundaries and T1/BOLD anatomy. For example, note that subject S2’s DN-B regions visibly do not overextend the amygdala in **Fig. 5**.

**Figure 5:**
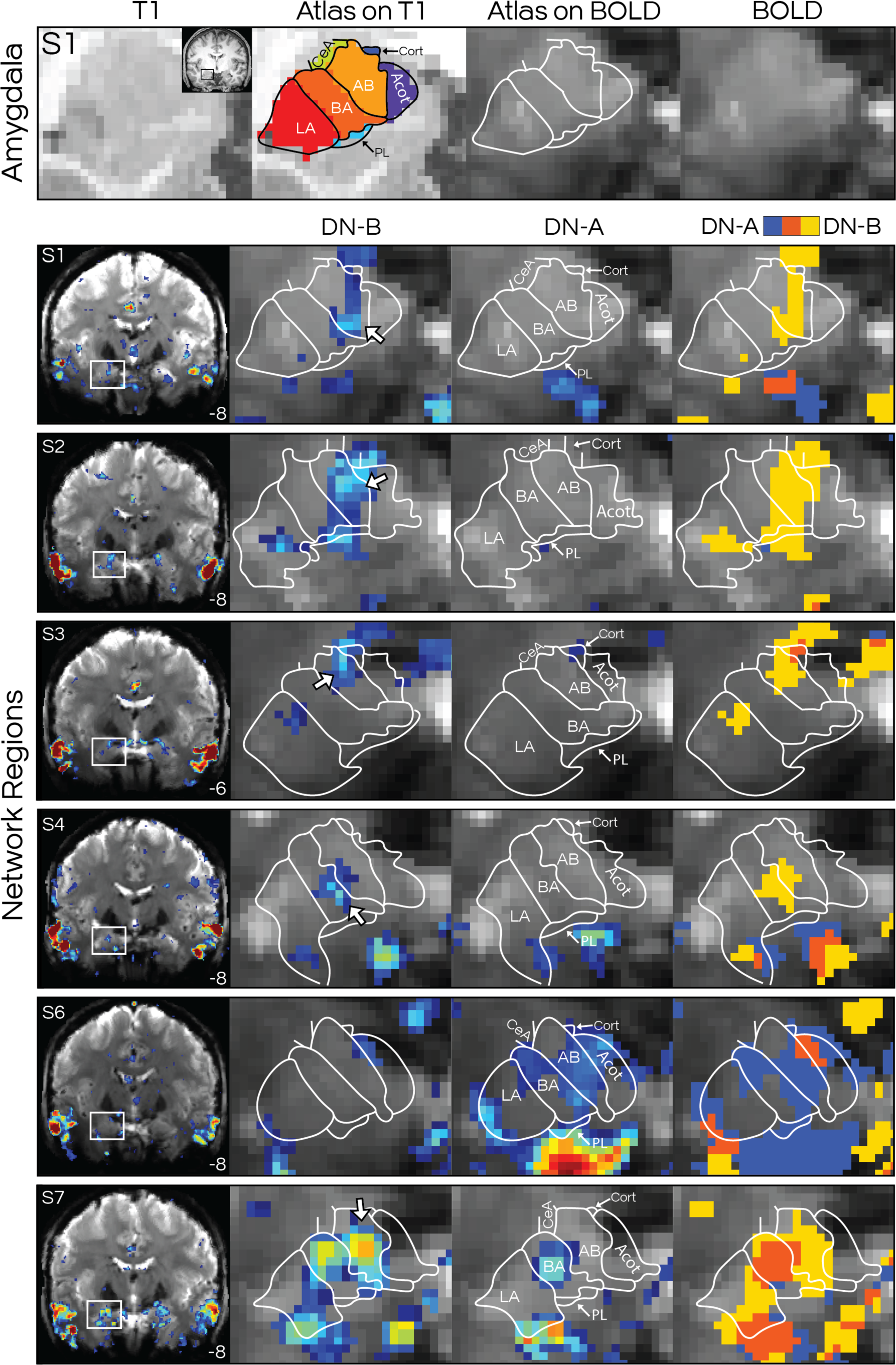
Individualized amygdala segmentation confirms regions of DN-B are located in or near the basolateral complex. Top row shows an example subject’s (S1) T1 and mean BOLD images, zoomed into the right amygdala, showing estimates of the amygdala nuclei from an automated, individualized segmentation.^121^ The remaining rows display the amygdala segmentation in each subject (S1–4, S6, S7), around MNI slice y = -6 to -9, along with the FC estimates of DN-A and DN-B, and their overlap. The left column shows the full coronal slice with the map of DN-B, with a box indicating the zoom-in location. White arrows indicate putative regions of DN-B which did not overlap with DN-A. In each subject, the regions of DN- B appeared to overlap most prominently with the accessory basal nucleus (AB) in each subject, often crossing the border into the basal nucleus (BA). Evidence for a distinct region can also be seen in ventral portions of the lateral nucleus (LA) in each subject. Further, DN-B regions also extended dorsomedially beyond the basolateral complex (see S1-S3), but notably did not overlap with the central nucleus (CeA). Other abbreviations refer to corticoamygdaloid (Acot), cortical (Cort), and paralaminar (PL) nuclei. **Supp. Fig. S8** shows the same views with the T1 as the underlay, to allow observation of each individual’s anatomy.

At a broad scale, the segmentation confirmed that the regions of DN-B were within the amygdala. When considering the nuclei demarcations, noteworthy consistencies were observed. In most subjects, DN-B reliably displayed: (1) a more prominent region at or near the accessory basal nucleus, sometimes extending to the adjacent basal nucleus (see arrows in **Fig. 5**), and (2) a separate, more lateral region within or near the lateral nucleus. These observations suggest that the DN-B regions were predominantly within the basolateral complex, which includes lateral, basal, and accessory basal nuclei.^123^ Finally, (3) DN-B regions often extended dorsomedially beyond the boundaries of the basolateral complex, towards the medial or ‘superficial’ aspect of the amygdala. We explore this dorsomedial region with targeted analyses in the next section.

To quantify these observations, we computed the spatial similarity of the DN-B map with the amygdala segmentation (**Supp. Fig. S4**). This analysis confirmed that DN-B regions overlapped most closely with the deeper, basolateral complex structures, and furthermore were predominantly in the accessory basal nucleus. These quantitative results were replicated and triplicated (**Supp. Fig. S4**). In contrast, when we repeated this analysis for DN-A, we did not observe the same pattern (**Supp. Fig. S4**). Hence, the individualized segmentation confirmed that, of the two interdigitated networks within the canonical DN, DN-B selectively includes regions in the amygdala, likely within the basolateral complex.

### Discovery of DN-B regions at or near the medial nucleus of the amygdala

In many individuals, DN-B regions extended beyond the dorsomedial boundaries of the basolateral complex, at or near where the central and medial nuclei are located. These protrusions strikingly did not overlap with the central nucleus, neither in the individualized (**Fig. 5**) nor group-defined segmentations^122^ (not shown). Comparison to the Mai atlas^124^ suggested these dorsomedial DN-B regions were at or near the medial nucleus (MeA), but also suggested that the automated parcellation had underemphasized (i.e., ascribed too few voxels to) the MeA. We therefore had an expert (author Q.Y.), who was blinded to the network maps, hand-draw the MeA in each participant in accordance with published procedures.^122^ In contrast to other amygdala nuclei, the medial nucleus can be more easily defined anatomically based on its location relative to the sulcal/septal anatomy and optic tract. The anatomical images used to delineate the MeA, alongside the hand-drawn estimates of the MeA, can be seen in **Supp. Fig. S8. Fig. 6** shows that in every participant, including subject S6, the manually drawn MeA either partially overlapped or bordered a region of DN-B. These medial regions of DN-B were located around MNI y = -3 to -5, and thus are likely distinct from the basolateral regions at y = -6 to -9.

**Figure 6:**
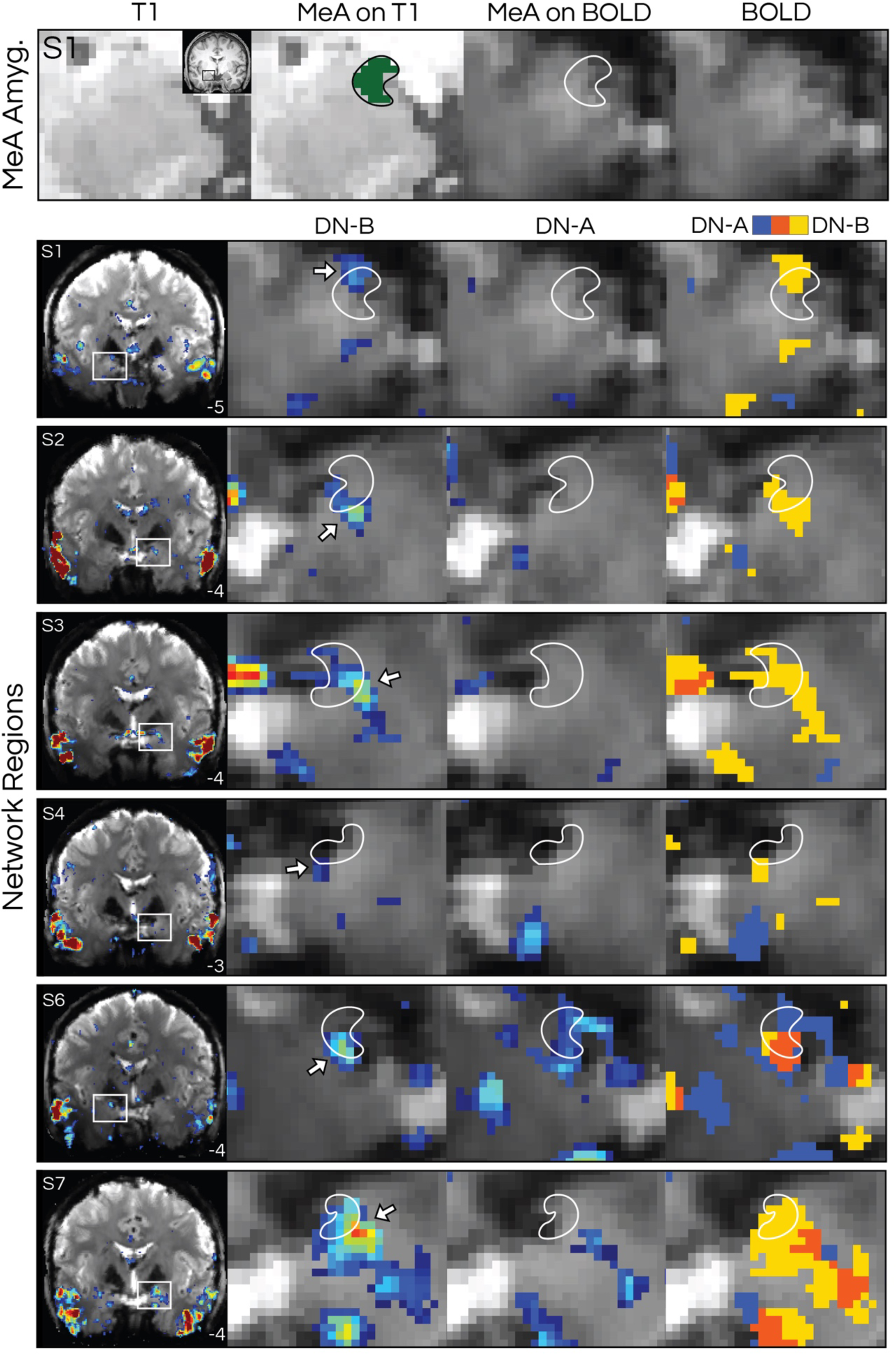
Hand-drawn segmentation confirms that DN-B regions are at or near the medial nucleus (MeA) of the amygdala. Figure formatted according to Fig. 5. The MeA was hand drawn by an expert blinded to the network maps, following Noto et al.^122^ and Mai et al.^124^ All subjects displayed a region of DN-B that partially overlapped or was adjacent to the estimate of the MeA, and which did not overlap with DN-A (see overlap map).

The regions were bilateral in 3/6 subjects and unilateral, on left or right, in the remaining 3 subjects (**Fig. 4**). We confirmed that this medial region was selectively connected to DN-B by placing a seed voxel near to the MeA in each subject. In all subjects, this analysis replicated the main components of DN-B (**Fig. 7**), often robustly. We also replicated and triplicated this medial amygdala region (**Supp. Fig. S7**). In two subjects (S4, S6) whose medial amygdala regions were unilateral in the discovery dataset, bilateral regions were found in the replication and/or triplication data. The MeA region was not present in subject S1’s replication dataset but was present in their triplication dataset. Conversely, in subject S4 a much clearer MeA region was detected in their replication compared to discovery dataset. These analyses provide reproducible evidence that network DN-B contains a region at or near the medial nucleus of the amygdala.

**Figure 7:**
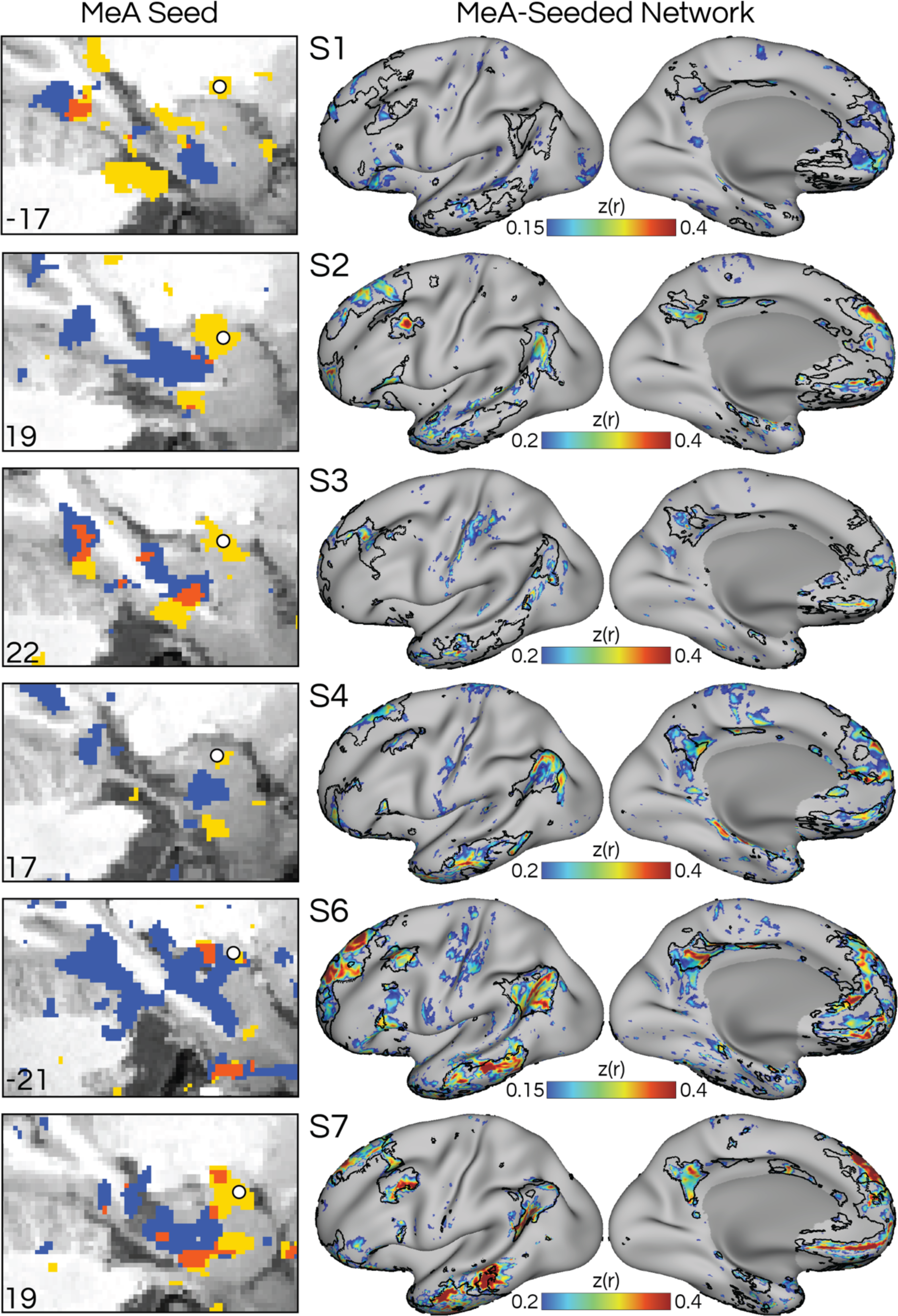
Seeds targeted to medial amygdala (MeA) selectively reproduce the distributed network DN-B. The left column shows a zoom-in on a sagittal view of the MTL showing each subject’s T1 and overlap map (similar to Fig. 3). FC maps (colorbar) from seeds targeting the MeA region of DN-B in each individual (white circles) recapitulated the distributed pattern of DN-B (see black outlines denoting surface-defined DN-B shown in Fig. 1), despite the amygdala showing reduced signal-to-noise ratio (**Supp. Fig. S1**). The strength of correlation varies across individuals (e.g., compare S7 with S1), which could be due to many factors, including data quality, signal dropout, the size of the amygdala region being targeted, and accuracy of seed selection.

### Confirmation of DN-B regions in the entorhinal cortex

Our volume-based analyses also revealed a clear region of DN-B at or near the entorhinal cortex (EC; **Fig. 3**), confirming a recent report.^86^ This region was observed in five out of six subjects (excluding subject S1 who had noticeably larger dropout in this region) and was confirmed by overlaying the correlation maps with hand-drawn MTL segmentations included in the NSD (not shown). In each subject, the entorhinal regions of DN-B were distinct from adjacent regions of DN-A. The slice locations where entorhinal regions were found ranged from MNI y = -7 to -14. The regions were also bilateral and replicated across datasets (not shown).

### Distributed association networks may be interdigitated along the hippocampal formation

In addition to the amygdala and entorhinal regions of DN-B, in two subjects (S2 and S4), we observed evidence that DN-B may include a region in the subiculum (see ‘s’ arrows in **Fig. 3** and seed ‘B’ in **Fig. 8**). This subiculum region of DN-B was adjacent to, but more posterior to a DN-A region in the subiculum that we previously described.^70^ This was an unexpected finding and raises the prospect that two distributed association networks, DN-A and DN-B, are finely intertwined along the long axis of the MTL. The DN-B subiculum regions were only clearly distinct from DN-A in subjects S2 and S4, and thus need to be replicated in follow-up studies.

**Figure 8:**
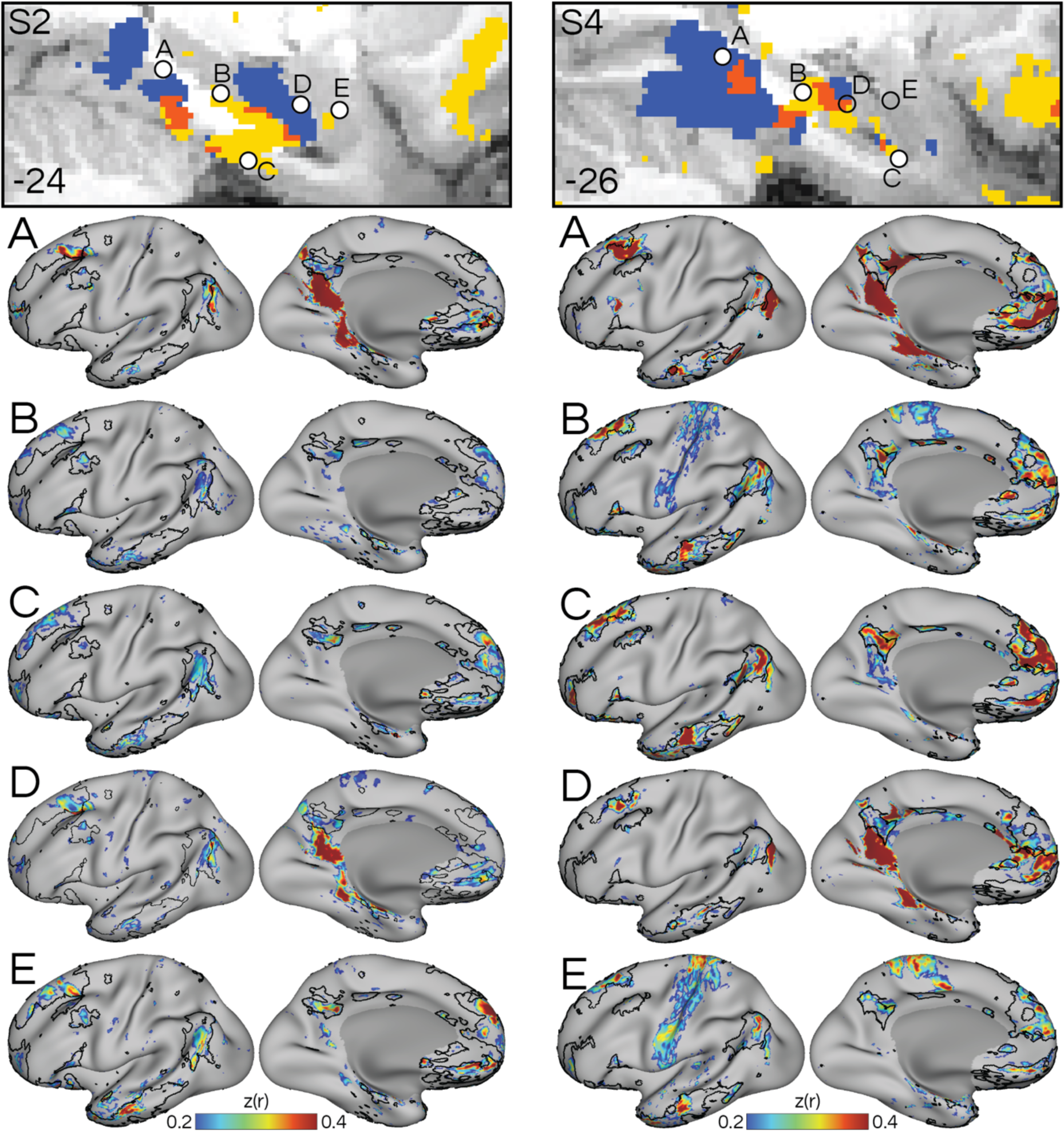
Two distributed cortical networks are interdigitated along the long axis of the medial temporal lobe (MTL). Circles indicate seeds that were hand-selected in two subjects (S2, S4) to target DN-A and DN-B along the MTL. Filled-in circles indicate seeds selected from the slice shown, and hollow circles indicate seeds selected in adjacent or nearby slices. Seeds targeted the (A) parahippocampal cortex (for DN-A), (B) subiculum (DN-B), (C) entorhinal cortex (DN-B), (D) subiculum (DN-A), and (E) basolateral amygdala (DN-B). Surface renderings underneath show the FC maps (colorbar) from the corresponding seeds, with a black border indicating network DN-B as defined in the surface (Fig. 1). In both individuals, seeds A and D reproduced network DN-A, while seeds B, C and E reproduced network DN-B, suggesting that DN-A and DN-B are interdigitated along the MTL. Note that the subiculum region of DN-B (seed B) was more posterior than the subiculum region of DN-A (seed D).

However, as an exploratory and hypothesis-generating analysis, we targeted seeds to each network along the long axis of the hippocampal formation in these two subjects. Moving from posterior to anterior MTL, **Fig. 8** shows that seeds targeting the two networks in parahippocampal cortex (DN-A), subiculum (DN-B), entorhinal cortex (DN-B), subiculum (DN- A), and the basolateral amygdala (DN-B) selectively and often strongly recapitulated the distributed organization of their associated networks (**Fig. 8**). This analysis emphasizes that the two networks can be defined from multiple parts of the MTL and suggests some level of interdigitation between the networks along the long axis.

## Discussion

This study investigated the connections of the MTL with distributed cortical association networks. We reproduced the observation that the canonical default network comprises two parallel distributed networks, DN-A and DN-B (**Fig. 1** and **Supp. Fig. S2**), and that only one, DN-B, is active during ToM (i.e., thinking about other people’s thoughts; **Fig. 2**). We discovered that DN-B contains bilateral regions within the amygdala (**Figs. 3 & 4**) that were previously missed,^20,66,70,86,115^ and reproduced these findings across (**Figs. 4–6**) and within participants (**Supp. Figs. S6 & S7**), and across analysis approaches (**Fig. 7** and **Supp. Fig. S3**). The high-resolution data allowed consideration of network regions in relation to individualized estimates of amygdala nuclei,^112,121^ which revealed that DN-B regions are located at or near the basolateral complex (**Fig. 5**), likely at or near the accessory basal nucleus (**Supp. Fig. S4**).

Potentially distinct regions were also reproducibly observed at or near the lateral nucleus (**Fig. 5**) and medial nucleus (MeA; **Fig. 6**), but not the central nucleus (**Fig. 5**). In contrast, DN-A did not show reliable evidence of containing amygdala regions (**Supp. Fig. S4**). The results support that the two networks, DN-A and DN-B, can be dissociated by their connections to posterior and anterior MTL. Finally, we observed evidence in 2 participants for closely juxtaposed, alternating regions of DN-A and DN-B along the long axis of the MTL (**Figs. 3 & 8**), suggesting the networks are interdigitated in the MTL, as they are in the cortex^66^ (**Supp. Fig. S2**). The results support that the amygdala and anterior MTL circuitry work with a broader distributed network of association regions (i.e., DN-B) to support socio-cognitive functions.

### Social cognitive regions of association cortex are intrinsically connected to the amygdala

Extensive research has described the circuits linking the amygdala, ventral and dorsal medial prefrontal cortex (including orbitofrontal cortex and anterior cingulate cortex), and temporal pole and their relevance to social cognition^110^ and psychiatric conditions.^125–132^ Here, we show that the amygdala-medial prefrontal circuitry is likely a part of a broader distributed association network that includes temporoparietal junction and dorsal posteromedial cortex, regions which are frequently implicated in ToM tasks^18,19,24,28^ and historically overlap the canonical DN.^21,22^ Previous human imaging has suggested a connection between the canonical DN (encompassing both DN-A and DN-B) and the amygdala^85,133–136^ (but see Roy et al.^84^; Sylvester et al.^85^; see also fifth figure in Bickart et al.^82^). Here, we show that it is specifically the portions of the canonical DN that are involved in social cognition (i.e., DN-B; **Fig. 2**) that are selectively connected to the amygdala. In contrast, the adjacent cortical network, DN-A, which is involved in mental scene construction (**Fig. 2**), did not show a connection to the amygdala (see overlap maps in **Fig. 5** and **Supp. Fig. S4**), and was instead confirmed to include posterior MTL regions (**Figs. 3, 7 & 8**). This distinction is notable as DN-A includes regions characteristic of the canonical DN, such as in the ventral posteromedial and retrosplenial cortex, yet here did not show a connection with the amygdala. We propose that these two distributed networks derive their dissociable functions from this differential link to the MTL; anterior MTL regions work selectively with DN-B regions to enable social cognitive functions, while posterior MTL regions work selectively with DN-A regions to enable episodic processes (**Fig. 2**).

The finding that association areas active during ToM are intrinsically connected to the amygdala, even while participants are at rest in the scanner, is notable. Evidence for amygdala activation during ToM tasks,^103–105,137,138^ suggests the amygdala is involved in ToM in an “online” capacity (but see Frith & Frith^1^). In contrast, evidence that only childhood lesions of the amygdala impair ToM^100,101^ sparked debate regarding whether the amygdala is only necessary during the acquisition of ToM abilities. Here, we show that in healthy adults the amygdala is intrinsically connected to the broader cortical network involved in ToM, supporting an online role. Indeed, our results suggest the connection is observable even in the absence of an active ToM task (i.e., at rest).

### DN-B regions are situated at or near the basolateral complex of the amygdala

We observed reproducible evidence (**Figs. 5 & 6** and **Supp. Fig. S4**) supporting that DN-B regions are located within the basolateral complex of the amygdala, which includes lateral, basal, and accessory basal nuclei.^123^ The basolateral complex is considered the main input structure to the amygdala, receiving widespread connections from the cortex.^126,139–142^ In contrast to other amygdala nuclei, the basolateral complex has expanded disproportionately in the mammalian lineage, likely due to the disproportionate expansion of neocortical areas providing its input.^10,143,144^ As DN-B is located deep within association zones that are most expanded in hominins,^2,5,145^ the presence of DN-B within the basolateral complex might have been predicted from evolutionary expansion patterns alone.

The basolateral complex is densely connected to the entorhinal cortex.^142,146^ Here, our results accord surprisingly well with this – albeit simplified – architecture: DN-B was shown to be connected to both the basolateral complex (**Fig. 5**) and entorhinal cortex (**Fig. 3**). The basolateral DN-B region overlapped most prominently with the accessory basal nucleus (**Supp. Fig. S4**), often extending into the basal nucleus (**Fig. 5**), and possibly includes a separate region within ventral portions of the lateral nucleus. Multisensory cortical input to the lateral nucleus is thought to be integrated through internal connections to basal and accessory basal nuclei, before being sent to the central and medial nuclei.^120,129,142,147^ The observation here of potentially distinct regions in different amygdala nuclei (**Figs. 5 & 6**) suggests that network DN- B may span multiple stages of the internal amygdala circuitry. Our results also support the conclusions of Aggleton et al.^148^ that amygdala projections from the orbitofrontal cortex, anterior insula, temporal pole, and anterior cingulate cortex – all of which contain regions of DN-B – predominantly target lateral and accessory basal nuclei (see also eleventh figure in Stefanacci & Amaral^147^; and sixteenth figure of McDonald^142^). Notably, in McDonald’s^142^ diagram, visual areas project to a more dorsal portion of the lateral nucleus, whereas ‘polysensory’ or associative projections are more ventral (**Fig. 5**). Thus, the specific patterns we observe here in the distribution of DN-B regions across amygdala nuclei appear to have precedent.

### DN-B regions are situated at or near the medial nucleus of the amygdala

The central (CeA) and medial (MeA) amygdala are evolutionarily older structures that mature early^149^ and originate from the subpallium (in contrast to the basolateral complex which is pallial).^120^ The CeA and MeA contain the main output projections to the autonomic system.^120^ The medial nucleus is closely linked to the accessory olfactory system in rodents, and projects abundantly to the hypothalamus.^150^ These projections are important for multiple olfactory-guided social behaviors in rodents, including mating, aggression, and parenting.^9,151–153^ Though few studies exist on the MeA in humans,^122^ it likely receives direct projections from the human olfactory bulb,^154^ suggesting conservation of function across species. Here, our blinded, hand-drawn estimate of the MeA partially overlapped with a DN-B region in every individual (**Figs. 6**). This region was bilateral (**Fig. 4 & Supp. Fig. S7**), was confirmed through seeding in the amygdala (**Fig. 7**), and was replicated and triplicated (**Supp. Fig. S7**). In contrast, we did not observe regions of DN-B in the CeA, neither using individualized (**Fig. 5**) nor group-defined (not shown) segmentations. Notably, the locations where we observed MeA regions of DN-B accord surprisingly well with where Brothers et al.^92^ identified neural responses when monkeys view socially relevant videos (see second figure in Brothers et al.^92^). Our findings suggest that the medial amygdala is also an important site for social cognitive functions in the human brain. Further supporting this link, in some individuals we detected putative regions of DN-B that may be at or near the ventromedial hypothalamus (see whole-brain images for S2, S3 and S4 in **Fig. 6**), which is the primary site of hypothalamic projections from the MeA in primates.^155^ Hence, although these results need to be confirmed in follow-up studies, the specific pattern of DN-B regions appears to recapitulate the proposed circuitry of the MeA from tracer studies.

### Networks for social and episodic functions are interdigitated along the medial temporal lobe

Our results support that there are anterior-posterior asymmetries in the network connections of the MTL. Although DN-A and DN-B are adjacent throughout the cortex, DN-B contains visibly larger regions in the anterior midline, whereas DN-A displays larger regions in the posterior midline (**Fig. 1** and **Supp. Fig. S2**). Our results here show that this asymmetry is also evident in the MTL when high-resolution mapping is performed. However, our results do not support a strict anterior-posterior division.^36,76,156^ Instead, both anterior and posterior MTL regions are connected to two parallel distributed networks that both include regions in posteromedial and anteromedial, as well as posterolateral and anterolateral cortex (**Fig. 1**).

Our present findings also accord with the idea that the long axis of the MTL differentiates spatial versus non-spatial (or contextual) information. Here, we show that the posterior MTL is more connected to a network involved in mental (spatial) scene construction (i.e., DN-A; **Fig. 2**), while the anterior MTL is connected to a separate network involved in social (non-spatial) cognitive processing (i.e., DN-B; **Fig. 2**). In two participants (S2 and S4), we also saw evidence that DN-A and DN-B are connected to adjacent regions of the subiculum (**Fig. 8**). Interestingly, the subiculum region of DN-B was located in a more posterior location than that of DN-A (i.e., compare seeds ‘B’ and ‘D’ in **Fig. 8**), a finding which also contradicts a strict anterior-posterior division of the MTL. These findings need to be confirmed in additional individuals. Speculatively, this fine-scale interdigitation might support the integrative model where contextual information from the anterior MTL, is integrated with spatial information in posterior hippocampus, via the subiculum.^157–160^

### Limitations and Technical Considerations

A limitation of the present study is that the distinction between networks relies on the correlation threshold chosen. Here, we focused on network regions that showed clear separation even at low correlation thresholds (0.15 and 0.20), and on findings that replicated across most individuals. The ideal threshold may vary across participants and datasets depending on data quality and smoothness. Here, we chose thresholds that were as low as possible, to limit noise (i.e., speckling) while still allowing regions of higher correlation (e.g., clear clusters of higher correlation) to be observed. Notably, within-network correlations were as high as 0.6 and above (e.g., note in **Fig. 3** how high-correlation regions are generally non-overlapping). Hence, the exact threshold used should not limit the interpretation of high- versus low-correlation regions, i.e., preferential connectivity to one network versus another. We previously noted that the degree of overlap between networks decreases as one moves from lower to higher resolution,^70^ indicating that overlap is a consequence of technical factors.

The replications of amygdala regions (**Supp. Figs. S6 & S7**) also showed minor variations. Sometimes, the replication and triplication datasets provided clearer evidence for amygdala regions (e.g., see subjects S3 and S4 in **Supp. Fig. S6**), and occasionally the replications failed (e.g., see subject S1’s replication data in **Supp. Fig. S7**). We have previously reported that even within individuals and on the cortical surface, where tSNR is higher, minor differences in FC maps are common when different datasets are analyzed.^71^ We believe these differences are due to small variations in registration (or data quality), meaning that even within individuals the same voxel might be sampling slightly different brain regions in different datasets. This highlights the difficulties of individualized network estimation when detailed anatomy is targeted. Here, we built confidence in our replications by visualizing the volume-defined networks on the surface, which confirmed that the same distributed networks were being targeted (**Supp. Fig. S5**).

## Conclusions

We show that in the human brain the distributed network associated with social cognitive functions is intrinsically connected to the anterior MTL and includes regions in the amygdala, entorhinal cortex, and potentially subiculum. We further show that the social cognitive network may contain multiple regions within the amygdala itself, putatively spanning different processing stages across the basolateral complex and medial nucleus, the latter of which is extensively linked to social behaviors.^9,92,151,152^ Our findings suggest that two networks within the canonical DN, DN-A and DN-B, which serve episodic and social functions, respectively, can be distinguished by their connectivity to the posterior and anterior MTL, respectively. These findings indicate that specific regions of association cortex derive their social cognitive functions through interactions with the amygdala and anterior MTL. We propose that network-targeted brain stimulation therapies for psychiatric and neurological conditions including autism, schizophrenia, and anxiety and major depressive disorder may be able to influence the amygdala by targeting stimulation to more accessible regions of DN-B in the cortex or cerebellum.^161,162^

## Acknowledgements

We thank Randy Buckner and Lauren DiNicola for helpful discussions. This work was supported in part by National Institute of Mental Health grant R00 MH117226 (R.M.B.) and an Alzheimer’s Disease Core Center grant (P30 AG013854) pilot award (R.M.B.) from the National Institute on Aging to Northwestern University, Chicago, Illinois.; a training award T32 NS047987 (to J.J.S. and N.A.); the William Orr Dingwall Foundations of Language Fellowship (J.J.S); and NSF GFRP award DGE-2234667 (D.E.). Collection of the NSD dataset was supported by NSF IIS- 1822683 and NSF IIS-1822929 (to K.K.). The content is solely the responsibility of the authors and does not necessarily represent the official views of the National Institutes of Health or National Science Foundation. This research was supported in part through the computational resources and staff contributions provided for the Quest high performance computing facility at Northwestern University which is jointly supported by the Office of the Provost, the Office for Research, and Northwestern University Information Technology, and the Center for Translational Imaging at Northwestern University.

## Author contributions

D.E., J.J.S., M.L., & R.M.B. designed the study; M.L., J.J.S., & R.M.B. collected the data; D.E., J.J.S., N.A., M.L., & Q.Y. analyzed the data with input from C.Z., K.K., & R.M.B.; D.E. & R.M.B. wrote the original draft, and JJ.S., N.A., M.L., Q.Y., C.Z., & K.K. edited and wrote the manuscript; R.M.B. secured funding.

## Methods

### Overview

Data processing and analysis procedures were previously reported in Kwon et al.^118^ and Allen et al.^111^

### Experimental model and study participants details

The study includes participants from two independent datasets. The Natural Scenes Dataset^111^ (NSD) and the Detailed Brain Network Organization (DBNO) dataset. The NSD is comprised of eight healthy human participants (2 males and 6 females; 19–32 years old) recruited from the University of Minnesota community. Sample size was chosen in accordance with the research team’s goal of collecting extensive data from a small number of individual subjects.^164^ Participants completed an initial screening session and 30–40 MRI sessions that included 2 resting-state fMRI scans alongside a visual image recognition task. The study ended with two task-based scan sessions measuring responses to other task contexts. This resulted in 33-43 MRI sessions per subject along with additional behavioral assessments. Participants provided informed written consent in compliance with procedures approved by the University of Minnesota Institutional Review Board and were paid for their participation.

The DBNO dataset includes 8 healthy human participants (4 females and 4 males; 22–36 years old). These participants were scanned at Northwestern University (downtown Chicago campus) and recruited from the local community. The sample size for this dataset was chosen based on past precision imaging work focused on estimating networks within extensively sampled individuals, while allowing replication across multiple individuals.^66,70^ Participants completed eight scanning sessions. During each session, participants provided two 7-min resting-state runs resulting in a total of 112 minutes (2 runs x 8 sessions x 7 min) of resting-state data per participant. In between the two resting-state runs in each session, participants completed nine active tasks targeting different cognitive domains. Participants provided written informed consent in compliance with procedures approved by Northwestern University Institutional Review Board and were paid for their participation.

### 7T fMRI data and quality control

Each participant was scanned across 30-40 sessions, providing an average of 2.0 hours of resting-state fMRI per participant. We performed rigorous quality control following a two-step process. First, head motion was estimated using FSL’s mcflirt command.^165^ Whole runs were automatically excluded if 1) maximum frame-wise displacement (FD) was greater than 0.4 mm, and 2) maximum absolute head motion (maxAbs) was greater than 2.0 mm. Second, any runs where FD exceeded 0.2 mm or maxAbs exceeded 1.0 mm were visually inspected for head motion and excluded if any could be clearly seen. Following quality control, two of the eight NSD participants were excluded due to excessive head motion. The remaining NSD participants retained 6-35 resting-state runs per participant (S1, 35 runs; S2, 6; S3, 16; S4, 12; S6, 19; S7, 18). For the two participants who provided 12-16 runs, half the data (6-8 runs) were assigned to a discovery dataset and the remaining half were assigned to validation dataset. For subjects with 18–19 runs, one-third of the data were assigned to discovery dataset and remaining two-thirds of the data were assigned to two validation datasets (6-7 runs each). For subject S1 who provided 35 runs, half of the data (17 runs) were assigned to a discovery dataset and the remaining half were assigned to two validation datasets (9 runs each). This is because we initially divided S1’s data into two datasets and began exploring half the data, before determining that stable estimates could be achieved with 6 runs.

### 7T MRI data acquisition and processing

As reported in Allen et al.^111^, 7T MRI data were collected using a 7T Siemens Magnetom scanner at the Center for Magnetic Resonance Research at the University of Minnesota. Blood-oxygenation-level-dependent (BOLD) images were collected using gradient-echo echo-planar imaging (EPI) at 1.8-mm isotropic resolution with whole-brain coverage with the following parameters: TR = 1,600 ms, TE = 22.0 ms, Flip angle 62°, FOV = 216 mm (FE) × 216 mm (PE), slice thickness 1.8 mm, slice gap 0 mm, matrix size 120 × 120, echo spacing 0.66 ms, bandwidth 1,736 Hz per pixel, partial Fourier 7/8, iPAT 2, multiband slice acceleration factor = 3, and 84 slices acquired in the axial plane. Dual-echo fieldmaps were collected periodically in each MRI session, and linearly interpolated over time for post-hoc correction of each EPI volume to account for interactions between small shifts in head position and field inhomogeneities (see further details in Allen et al.^111^). The accuracy of this fieldmap distortion correction was confirmed through comparison to the magnitude image, which is not distorted.

Initial pre-processing as part of the NSD release includes slice timing correction, alignment of each volume to correct for head motion within a run, alignment of data across sessions, and correction for EPI distortion. Registration steps were all performed in one interpolation to reduce smoothing. Functional images were co-registered to the T1 image using a non-linear warp implemented in ANTs 2.1.0^166^ (BSplineSyN with parameters [0.1, 400, 0, 3]). This transformation was visually confirmed to be accurate based on comparisons to the original T1 volume, as outlined in detail in Allen et al.^111^

For functional connectivity analysis, we performed additional preprocessing on the resting-state data following the steps outlined in the iProc pipeline.^70^ The first ten volumes (approximately 12 seconds) were removed to accommodate for T1 attenuation effects, and an individual-specific mean BOLD template was generated from included runs. There is a slight flaring of the brain image at the superior aspect in the meanBOLD image (See **Fig. 3**). This flaring is a result of fieldmap measurement inaccuracies outside the brain and does not affect measurements within the brain. Nuisance variables, including six parameters to account for head motion, as well as whole-brain, ventricular, and deep white matter signal, and their temporal derivatives, were calculated and regressed out of the data. Nuisance regression was performed using 3dTproject (AFNI version 2016.09.04.1341^167^) on the native-space-projected BOLD data resampled to 1mm isotropic resolution (i.e., the ‘func1p0mm’ version of the NSD data). Data were then bandpass filtered at 0.01-0.1Hz (using 3dBandpass from AFNI). After pre-processing, we utilized a temporal signal to noise ratio (tSNR) map to examine data quality in the MTL. We found that there was little to no dropout in MTL regions, namely the amygdala, as the majority of dropout observed was in more ventral regions of the temporal lobe (See **Supp. Fig. S1**).

### Surface-based functional connectivity analysis at 7T

For surface-based analysis, preprocessed data were projected onto a standardized cortical surface containing 163,842 vertices (fsaverage) per hemisphere using FreeSurfer’s vol2surf command^112^ and smoothed along the surface using a 2.5mm full-width at half-minimum (FWHM) Gaussian kernel. The highest-resolution fsaverage cortical mesh was used to minimize blurring and preserve fine-scale distinctions between networks. The smoothing kernel was chosen based on preliminary analyses in one individual to optimize the trade-off between minimizing smoothing, maximizing correlation values, and minimizing noise or ‘speckling’ in the correlation maps. Functional connectivity matrices were calculated in each participant by computing vertex-vertex Pearson’s product-moment correlations for each run, r-to-z transforming, and then averaging across runs within each dataset. These matrices were used for surface-based network estimation using an interactive platform for manual seed-selection and correlation map viewing based in Connectome Workbench.^168^

Our initial analyses focused on identification of DN-A and DN-B based on characteristic anatomical distinctions between the networks reported in Braga & Buckner^66^ and Braga et al.^70^, including the replicated distinction between the networks in posteromedial cortex, inferior parietal lobe, and lateral temporal cortex. Briefly, we aimed to identify in each individual a network that follows the expected pattern for DN-A of containing regions in ventral PMC and/or retrosplenial cortex, posterior parahippocampal cortex, and posterior inferior parietal lobe. We also aimed to identify in each individual a network that followed the expected pattern for DN-B, which occupies adjacent, but distinct regions to DN-A in multiple cortical zones. Specifically, DN-B contains regions within more dorsal PMC at or near the posterior cingulate cortex, and more rostral inferior parietal regions at or near the temporoparietal junction. DN-B also contains prominent regions in lateral temporal cortex and inferior frontal cortex. To define the networks, we used two different approaches: a manual seed-based and a data-driven clustering or ‘parcellation’ approach. In the seed-based approach, initial seed selection targeted the dorsal lateral prefrontal cortex to (i) align with our previous studies and (ii) allow characteristic patterns of functional connectivity to be appreciated in distal zones (e.g., in the MTL and posteromedial cortex) without the confound of local blurring near the seed. To limit the effects of observer bias in the manual seed-selection process, we also employed a multi-session hierarchical Bayesian model (MS-HBM) parcellation method.^114^ In each individual, we used a *k* value (i.e., number of clusters) between 14-20 and selected the solution that best matched the networks observed by the seed-based approach (*k* = 14). The clustering analysis provides a data-driven confirmation of the patterns observed in the manual seed-based approach, offering converging evidence for the distinction between DN-A and DN-B.

### Volume-based functional connectivity analysis at 7T

To examine regions of DN-A and DN-B in the MTL, we estimated the networks using a seed-based approach in the volume. We analyzed native-space projected volumetric BOLD data from the NSD that was preprocessed for functional connectivity analysis prior to surface projection.

Based on data quality and the strength of correlation maps observed in initial explorations, a 2.5mm FWHM smoothing kernel was applied to five subjects (S1-S4,S6) and a 2mm FWHM smoothing kernel was applied to one subject (S7) using fslmaths (FSL v6.0.3).^165^ Data were analyzed and visualized using AFNI’s InstaCorr.^167,169^ 3dSetUpGroupInCorr was used to calculate a Pearson’s product-moment correlation coefficient between all voxel pairs within a whole-brain brain mask for each run of resting-state data. Each matrix was Fisher transformed prior to cross-run averaging with 3dGroupInCorr to create a single, cross-run average functional connectivity matrix for each dataset from each individual. We used AFNI to interactively select individual voxels as seeds and observe their associated whole-brain correlation maps. This process was used to define DN-A and DN-B in the discovery dataset. Initially, we converted the surface-selected seed vertex locations that defined DN-A and DN-B into approximate X, Y, Z voxel indices to define the same networks in the volume. Perhaps because the surface projection applies an interpolation which means that a surface vertex may not correspond directly to a single voxel, these converted seed locations did not produce maps with the expected separation of DN-A and DN-B as seen in our prior work.^70^ We therefore manually explored the surrounding voxels for a seed that better distinguished the two networks. Seed optimization was performed by consulting cortical correlation patterns and focusing on the anatomy of regions within the posteromedial cortex. We explicitly did not consult the MTL during seed selection, and selected candidate DN-A and DN-B seeds based on their degree of overlap with surface-defined maps from the data-driven clustering analysis (e.g., see **Figs, 1, 7 & 8**).

Overlap maps were created by thresholding the correlation maps at z(r) > 0.3 and then binarizing each network separately. The DN-B binarized map was multiplied by a scalar value of 2, from which we then subtracted from the binarized DN-A map. This created an image where network DN-A’s voxels had a value of -1, network DN-B’s voxels had a value of +2 and the overlap voxels had a value of +1. The overlap maps were visualized in wb_view^168^ using the PSYCH color bar set to range from -2.5 to 0 and 0 to +20.

To aid the detection of network regions in consistent anatomical location across subjects, we projected the volume-defined functional connectivity maps, mean-BOLD, and T1 images to MNI152 space. Projection was done using the nsd_mapdata commands provided with the NSD, which applied precomputed transforms using nearest neighbor interpolation.^111^

### Replication and Triplication at 7T

To ensure robustness of findings, after analysis of the discovery datasets the results were replicated and triplicated within the same individuals. Initially, the same seed vertices used to define the networks in the discovery dataset were used to define the networks in the left-out datasets. This broadly replicated the definition of the same networks, but sometimes reduced the differentiation between DN-A and DN-B compared to what was seen in the discovery dataset, or in our previous work at 7T.^70^ Hence, we optimized the seeds in the validation datasets by searching for a nearby seed voxel that maximized correlations within characteristic DN-B regions and minimized correlations in DN-A regions. Again, we explicitly did not consult the MTL in the seed selection procedure, and based seed selection on the distinction between the networks observed in the posteromedial cortex, as well as with the degree of overlap with surface-defined networks from the data-driven clustering analysis. We have previously noted that the optimal seed location to target a given network can change slightly across datasets, even from the same individual, potentially due to small registration differences across datasets.

### Determining overlap with amygdala nuclei

To navigate the anatomy of the MTL and quantify overlap of network regions with specific anatomical regions, we utilized (i) a previously published hand-drawn atlas of amygdala subregions in group-averaged MNI152 space,^122^ (ii) hand-drawn maps of MTL regions that are included in the NSD and were defined within each individual (but do not include the amygdala), an automated Amygdala segmentation that was calculated using FreeSurfer (7.3.2),^121^ and a hand-drawn estimate of the medial nucleus by an expert observer (author Q.Y.) who was blind to the network maps. The FreeSurfer tool uses a procedure where a segmentation of the amygdala into 9 nuclei, defined from super-high-resolution (100-150 µm) 7T imaging in 10 post-mortem specimens, is template-matched to anatomical images from each individual. The tool therefore provided the location of 9 amygdala nuclei in native space T1, including the accessory basal (AB), basal (BA), lateral (LA), medial (fs-MeA), central (CeA), cortical (Cort), cortico- amygdaloid-transition (Acot), anterior amygdaloid area (Ant), and paralaminar (PL). Atlas regions were projected into MNI space for visualization alongside DN-B and DN-A regions using nsdmapdata^111^ and fslswapdim (FSL).^165^ The resulting parcellation was cross-checked with the Mai atlas^124^ for accuracy.

### Manual delineation of the medial amygdala nucleus

Because the FreeSurfer amygdala parcellation ascribed few voxels in each individual to the medial nucleus (MeA), we had an expert observer hand-draw the MeA bilaterally in each individual based on procedures and anatomical landmarks reported in Noto et al.^122^ and Mai et al.^124^ This observer (author Q.Y.) was blinded to the network maps. The MeA was drawn based on the anatomy captured by each participant’s T1 image. The T1 image projected to 1-mm MNI space was used, to adhere to Noto et al.^122^ The anterior limit of MeA coincides with the posterior edge of temporal piriform cortex,^124^ typically around MNI coordinate y = -4. First, the posterior limit of piriform cortex was identified,^170^ and the anterior edge of the MeA was marked on the next coronal slice, 1.0 mm in the posterior direction from the posterior edge of piriform cortex. In this coronal slice, the surface of the entorhinal sulcus changes from flat and smooth when contiguous with piriform cortex, to angled and exhibiting a distinct small bump, which indicates the location of the cortical amygdala. This bump is visible in conventional T1 images. Based on Mai et al.^124^, from this anterior edge, the MeA region of interest was extended 3-4 mm into the amygdala for the posterior border of the MeA. The posterior border of the MeA ROI was identified using the optic nerve as a landmark. Specifically, the MeA continues in the posterior direction until the optic nerve nearly touches the amygdala surface, and the anterior border of the hippocampus becomes visible. In the lateral extent, the MeA extends 4mm into the amygdala based on Mai et al.^124^

### Quantifying overlap with amygdala nuclei

To quantify the overlap between the network regions and the amygdala segmentation, in each subject, we restricted the analysis to a mask based on the freesurfer parcellation that included the whole amygdala. Within this mask, we binarized the thresholded correlation maps, using our previously defined thresholds (**Fig. 4**). We then used the dice similarity function in MATLAB to compare spatial overlap between the binarized correlation maps and each atlas region (i.e., nuclei). This was repeated for the replication and triplication datasets to provide up to three independent estimates of the overlap in each participant. We also assessed overlap with the group-average atlas of the amygdala, obtained from Noto et al.^122^, which broadly replicated that DN-B regions were at or near the basolateral complex and medial nucleus, but not central nucleus (not shown). The correspondence observed using the group-average amygdala atlas was not as clear as the individualized maps, which showed consistent relationships across participants (**Supp. Fig. S4**).

### Replication of network distinction and functional dissociation at 3T

Analysis of resting-state data from the Detailed Brain Network Organization (DBNO) dataset was previously reported in Kwon et al.^118^ Eight individuals were scanned at Northwestern University. Participants were native English speakers, neurologically healthy, and had normal or corrected-to-normal vision (4 females, age range 22-36 years, mean age = 26.8). Participants provided written informed consent in compliance with procedures approved by Northwestern University Institutional Review Board and were paid for their participation. The experiment consisted of eight MRI sessions, collected over approximately 18 - 63 days, based on participant availability and avoiding consecutive days. During each session, participants completed two 7-min resting-state runs resulting in a total of 112 minutes (2 runs x 8 sessions x 7 min) of resting-state data per participant. During the resting state runs, participants were asked to fixate their eyes on a cross at the center of the screen. The position of the screen was adjusted for each participant and session to ensure a comfortable viewing angle to minimize head motion. The resting-state runs were collected as the first and last runs in each session.

### Acquisition, processing, and quality control of 3T MRI data

Functional images were collected at the Center for Translational Imaging at Northwestern University on a 3T Siemens Prisma scanner. A high-resolution T1-weighted magnetization-prepared rapid acquisition gradient echo (TR = 2,100 ms, TE = 2.9 ms, FOV = 256 mm, flip angle = 8°, slice thickness = 1 mm, 176 sagittal slices parallel to the AC-PC line) was acquired after the first resting-state run. Multi-echo BOLD data^171,172^ were collected using a 64-channel head coil with the following parameters: TR = 1,355 ms, TE = 12.80 ms, 32.39 ms, 51.98 ms, 71.57 ms, and 91.16 ms, flip angle = 64°, voxel size = 2.4 mm, FOV = 216 mm x 216 mm, slice thickness = 2.4 mm, multiband slice acceleration factor = 6.

Functional MRI data were pre-processed using the iProc pipeline^70^ with changes implemented to accommodate multi-echo data described below. The iProc pipeline is optimized for alignment of data from different sessions and runs within each individual, using individualized registration templates and with registration steps performed in a single interpolation to minimize blurring and preserve anatomical details. Runs with excessive head motion (a maximum framewise displacement > 0.2 mm or a maximum absolute displacement > 1 mm) were visually inspected and excluded if they exhibited clear movement. This resulted in a total of 10-16 resting-state runs per participant (S1: 16; S2: 16; S3: 16; S4: 6; S5: 10; S6: 15, S7: 14; and S8: 14 runs). For the seven participants providing 12 or more runs, the data were divided into two groups: a discovery dataset (7-8 runs) and a validation dataset (7-8 runs). The DBNO analyses focused on the discovery dataset only.

The first nine volumes (approximately 12 seconds) were removed to account for the T1 attenuation artifact and a mean BOLD template was generated for each individual using the included runs (see details in Braga et al.^70^). Functional data were registered to a 1.0-mm isotropic resolution T1 image for each individual via this mean BOLD template, including within-run motion correction and across-run and -session alignment. Brain extraction was performed on the T1 using FSL’s Brain Extraction Tool (FSL v6.0.3). White matter and cerebrospinal fluid masks were registered to the skull-stripped mean BOLD template to calculate mean white matter, cerebrospinal fluid, and whole brain signal time series. These mean time series, along with 6 head motion parameters and temporal derivatives, were used to remove nuisance signals through regression. Data were bandpass filtered at 0.01-0.10 Hz. Preprocessed data were then projected onto a standardized cortical surface (fsaverage6: 40,962 vertices per hemisphere) using Freesurfer^112^ and smoothed along the surface using a 2mm FWHM Gaussian kernel.

### Data collection for social cognitive tasks

In each session, DBNO participants completed nine active tasks targeting different cognitive domains. Two tasks targeted social cognitive or ‘theory of mind’ (ToM) processes: False Belief and Emotional Pain,^15,16^ and were previously described in DiNicola et al.^20^ Each task lasted 319 seconds per run, with 4 runs of each task collected per subject. Each run included 10 trials. The False Belief task required participants to make inferences about another person’s beliefs in either social (false belief condition) or visual (false photo condition) contexts.^15^ For the false belief condition, participants responded true or false to prompts, such as: “Lillian titled her novel How to Cook a Grouse and submitted it. Her publisher retitled the novel How to Cook a Pheasant. In the bookstore, Lillian notices the title is How to Cook a Grouse, True or False?” For the false photo condition, participants responded to true or false questions, such as: “A woman’s drawing of her back yard included a blue treehouse. That was before the storm. She built a new treehouse last summer and painted it red. Today the treehouse is blue, True or False?” The Emotional Pain task required participants to provide a rating for another person’s emotional or physical pain. To rate emotional pain participants responded to prompts such as: “Alison lives by herself. Every week her son comes to visit her and they go out to lunch. Her son just lost his job and is arguing with his wife. One day Alison gets a call from her son who tells her that he has been diagnosed with cancer. Rate protagonist pain or suffering.”^16^ To rate physical pain participants responded to prompts such as: “Suzie was riding in a cab. When she opened the door and began to step out, a child accidentally bumped the door and it close on Suzie, smashing her leg.” Participants were trained on the tasks outside the scanner prior to the first session and were given an example trial in the scanner prior to the start of the first run in each session, to accustom them to the pace of the task. For both tasks, each run started with 18s of fixation to a centrally presented crosshair (‘+’), and each trial included 10s of story presentation, and a 5-second questions presentation. There was 15 seconds of fixation in between each trial. Trial and block order was counterbalanced within and across runs (see DiNicola et al.^65^).

### Data collection for mental scene construction task

In each DBNO session, participants also completed a mental imagery task. Participants were asked to covertly imagine scenarios based on a specific prompt provided on a screen. Each run comprised 25 trials. In each trial, participants saw a written description of a scenario, such as “Imagine a nature trail in the summer,” and were instructed to imagine the scenario in their mind. The imagination period lasted 7s. After the imagining period, participants were given three seconds to rate the visual vividness of their imagined scenario on a four-point scale (“Nothing”, “Vague”, “Moderate”, and “Vivid”), followed by an additional three seconds to rate the auditory vividness of their imagining on the same scale. In half of the runs, the auditory vividness rating preceded the visual vividness rating. After both ratings, a fixation cross was presented for 10-11 seconds (jittered), followed by the next trial. Each run began with 18s of fixation to a crosshair (‘+’), and the mental imagery task was always the second-to-last task of the session (followed by a final resting-state run). Each run lasted 614 seconds and eight runs were collected from each subject.

After the MRI scanning session, participants immediately completed an out-of-scanner follow-up task on a computer. In this task, participants were re-presented with the 25 scenarios they had seen during the in-scanner mental imagery task. For each scenario, participants were asked to provide ratings to questions regarding their experience imagining inside the scanner. First, participants repeated their visual and auditory vividness ratings (using the same scale), which confirmed that participants were able to replicate their in-scanner responses in the out-of-scanner test (correlation between in- and out-of-scanner vividness ratings: S1, 0.81; S2, 0.93; S3, 0.81; S4, 0.79; S5, 0.69; S6, 0.92; S7, 0.61; S8, 0.7). Participants then provided ratings on 15 additional questions using a 9-point endorsement scale (ranging from Strongly Disagree to Strongly Agree). These questions probed additional aspects of the imaging experience such as the degree to which they thought about spatial dimensions (i.e., “I envisioned the location of objects, people or places”), their level of immersion (i.e., “I felt immersed in my thoughts, as if I was experiencing the item(s) in real life”), and how successful they were at imagining the scenario (i.e., “I was able to imagine what was asked successfully”). Participants were given unlimited time to provide their responses for each scenario and question, and on average spent 21.9 minutes on each follow-up session (range: 12.3 - 48.6 minutes).

### Resting-State Functional Connectivity Analysis at 3T

Within each surface-projected resting-state run, vertex-wise Pearson’s product-moment correlations were computed to generate a correlation matrix. Correlation matrices were r-to-z transformed before averaging across runs within each participant. We defined DN-A and DN-B in the 3T data using the same process as described in the 7T data, including seed-based and data-driven clustering approaches. The MS-HBM parcellation method was again applied to this dataset to define networks,^163^ which is supposed to improve stability in the network estimation procedure. In each individual, we used a *k* value between 14-20 and selected the solution (*k* = 14) that best matched the networks observed by the seed-based approach.

### Task-Based Analysis at 3T

Task data were quality controlled for head motion and tSNR using the same thresholds and procedures used for the resting-state data. Runs that passed quality control for MRI data were then assessed for behavioral performance. Behavioral performance was computed for each participant and task. Error rates include invalid responses (e.g., multiple button presses), non-responses, and incorrect responses. For the False Belief task, runs with <80% accuracy were excluded from further analysis resulting in the following remaining runs for each subject: S1, 4 runs; S2, 4; S3, 2; S4, 0; S5, 4; S6, 3; S7, 1. Error rates across the remaining runs were S1, 7.5%; S2, 5.0%; S3, 0.0%; S4, n/a; S5, 10.0%; S6, 6.7%; S7, 15.0%; S8, 0.0%). For the Pain

Task, which involved a subjective rating, error rates included invalid and non-responses, and were as follows: S1, 0%; S2, 0%; S3, 2.5%; S4, 0%; S5, 0%; S6, 0%; S7, 2.5%; S8, 0%). No runs were excluded from the Pain task. An initial analysis that included all runs across all subjects (i.e., including runs with high error rate) produced very similar results to those in **Fig. 2**.

For the mental imagery task, after exclusion due to head motion and tSNR criteria each participant retained 6-8 out of 8 runs (S1, 8 runs; S2, 8; S3, 6; S4, 8; S5, 6; S7, 6; S8, 7). As this task was analyzed in a trial-wise manner, error trials (missed or invalid responses) were excluded trial-wise. Participants on average provided both visual and vividness ratings for 96.4% of trials per run (S1, 100%; S2, 100%; S3, 100%; S4, 92%; S5, 95%; S6, 100%; S7, 86%; S8, 97%). Any trial missing at least one response was excluded from analyses, but was still modelled in FEAT.

Task BOLD data were analyzed using a general linear model (GLM) as implemented in FSL’s FEAT.^173^ For the ToM tasks, each condition was modelled as a single explanatory variable that included both the story and question periods of each trial. For the mental imagery task, each trial element was modeled individually, with separate explanatory variables for each trial’s imagining, response, and post-response fixation periods. Explanatory variables were convolved with a double-gamma hemodynamic response function. In the False Belief task, the False Belief condition was contrasted against the False Photo condition. In the Pain task, the Emotional Pain condition was contrasted against the Physical Pain condition. In the mental imagery task, each imagining period was contrasted against its preceding fixation period to create a beta estimate for each trial.

Task contrast maps for the ToM tasks were converted to t-statistics and then to a z-statistic, then averaged together across runs within each participant using fslmaths.^165^ Next, we averaged together the contrast maps from the False Belief and Emotional Pain tasks using fslmaths (FSL v6.0.3)^165^ to create a single map of ToM-related activity per subject. We overlaid this map against the outline of the functional connectivity-defined estimate of DN-B to assess the extent of overlap between the two maps. The close correspondence between maps (**Fig. 2**) replicated the findings of DiNicola et al.^20^ that DN-B, as defined using functional connectivity, captures brain regions involved in social cognitive processes.

For the mental imagery task, a behavioral summed composite metric was created for each trial based on the participant’s responses to the visual vividness (in-scanner) and spatial (out-of-scanner) ratings. These closely align with the “Scene Construction” composite reported in DiNicola et al.^65^ using a similar approach. For each subject, trials were identified with the highest and lowest Scene Construction scores (top 20% of usable trials and bottom 20% of usable trials; if multiple trials at the cut-off margins had the same score, we randomly selected which were included. Beta values for trials in the bottom 20% of composite scores were multiplied by -1, and then all trials (top and bottom 20%) were averaged together to create a contrast map between high-Scene Construction and low-Scene Construction trials. The contrast (beta) map was then z-scored to create a z-stat map. We compared this map to the functional connectivity-defined estimate of DN-A to determine how Scene Construction-related activity overlaps with the anatomy of DN-A. The close correspondence between maps (**Fig. 2**) confirms that DN-A, as defined using functional connectivity, can delineate brain regions involved in mental scene construction (i.e. “episodic projection” or EP).

---

**Supplementary Figure S1:**
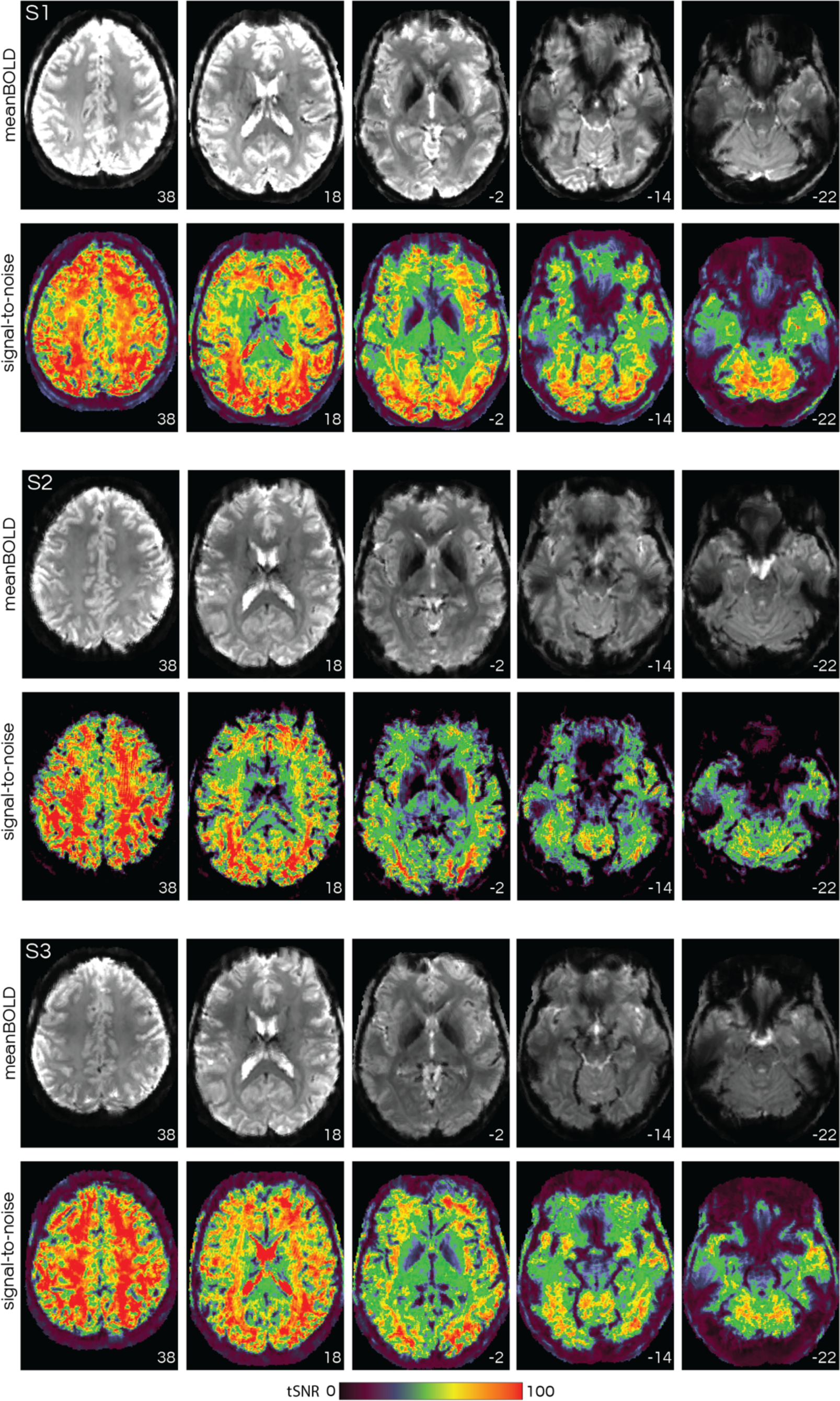
Average blood-oxygenation-level-dependent (BOLD) and temporal signal-to-noise ratio (tSNR) images for 3 example participants (S1–S3) display the high quality of the high-resolution 7T data, including in the anterior medial temporal lobe. Numbers in the bottom right correspond to MNI coordinates for each axial slice. Left hemisphere is on the left of each view.

**Supplementary Figure S2:**
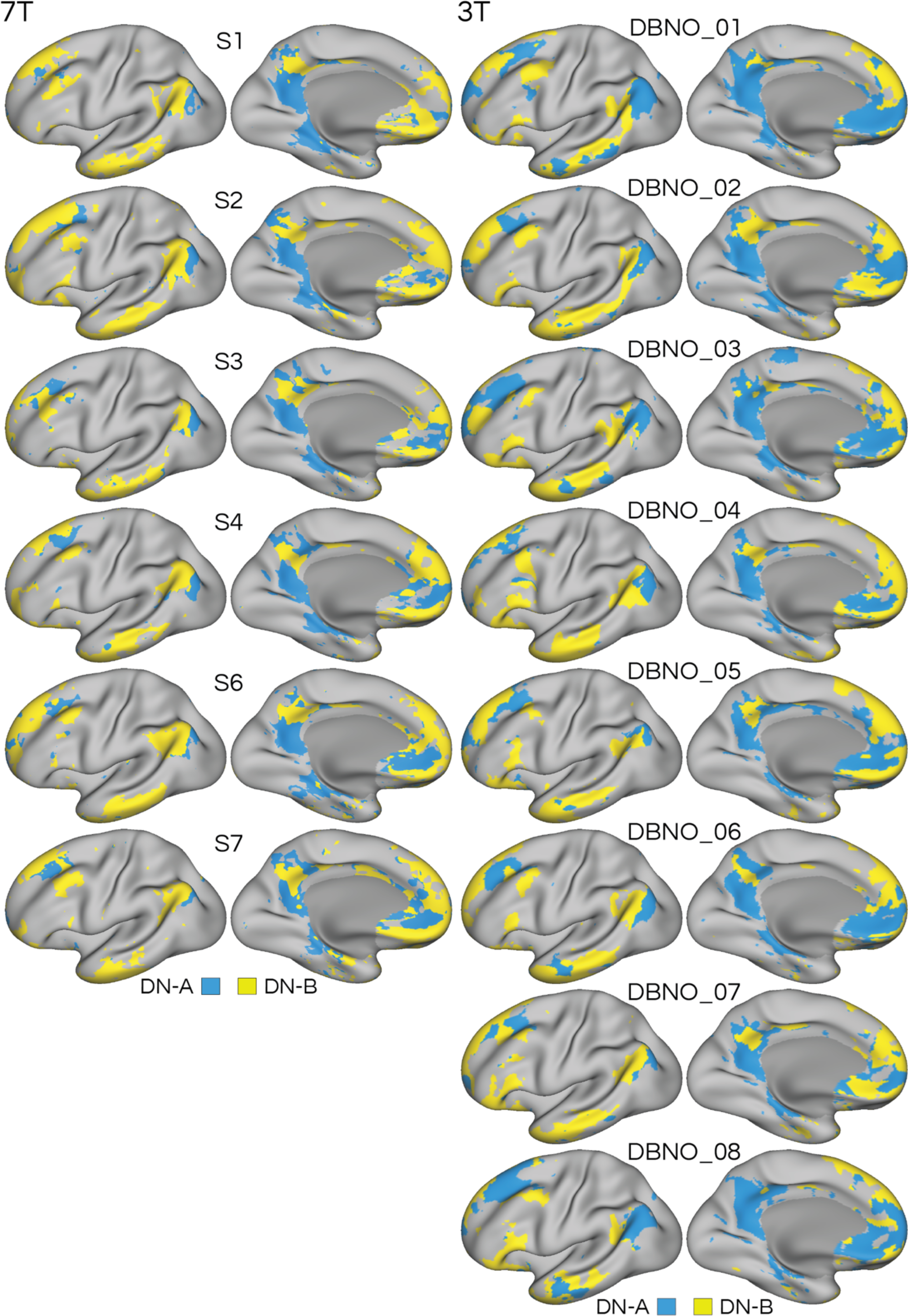
Replication of default networks A (DN-A) and B (DN-B) using functional connectivity at 3T in 8 individuals from the detailed brain network organization (DBNO) dataset. Network estimates were made using data-driven clustering of functional connectivity patterns within each individual, using the same approach as used in the 7T data. Comparison of the 7T and 3T data confirms that the same two networks were targeted in each subject and dataset, despite small variations in the shape and location of regions which are expected given idiosyncrasies of each individual’s brain. Note how DN-A (blue) contains characteristic regions in the ventral posteromedial cortex (PMC) and posterior inferior parietal lobe (IPL), as well as the posterior parahippocampal cortex, in each individual. Similarly, DN-B (yellow) contains a more dorsal PMC region, at or near the posterior cingulate cortex, as well as a more anterior region of IPL at or near the temporoparietal junction (TPJ). Differences were also present between the datasets: note how the 7T data revealed evidence for a region of DN-B in the anterior medial temporal lobe, whereas the 3T data did not. For DBNO participants, data from ToM and mental imagery tasks were used to replicate the functional dissociation of the two networks and confirm that network DN-B encompasses regions active during a social cognition task (Fig. 2).

**Supplementary Figure S3:**
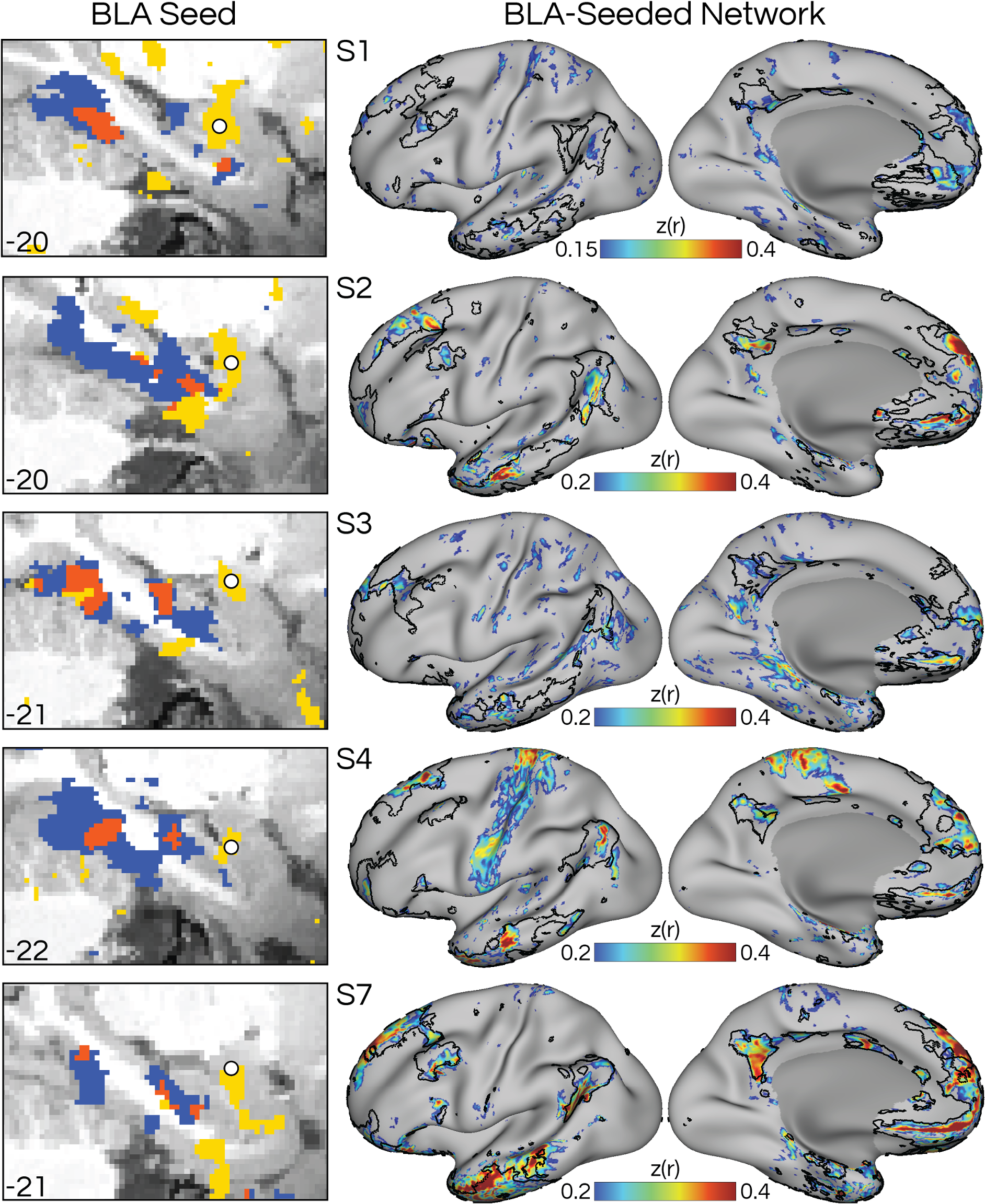
Seeds targeted to the basolateral complex of the amygdala (BLA) reproduce the full distributed network DN-B, despite the low signal-to-noise ratio in these regions, confirming that DN-B contains a region in the amygdala. The left column shows a zoom in on the MTL on a sagittal slice in five subjects (S1-4, S7) where a distinct region of DN-B can be seen (Fig. 5). Figure formatted according to Fig. 7. The strength of correlation varied across individuals (e.g., compare S7 with S1), which could be a result of many factors, including data quality, signal dropout, the size of the amygdala region being targeted, and accuracy of seed location.

**Supplementary Figure S4:**
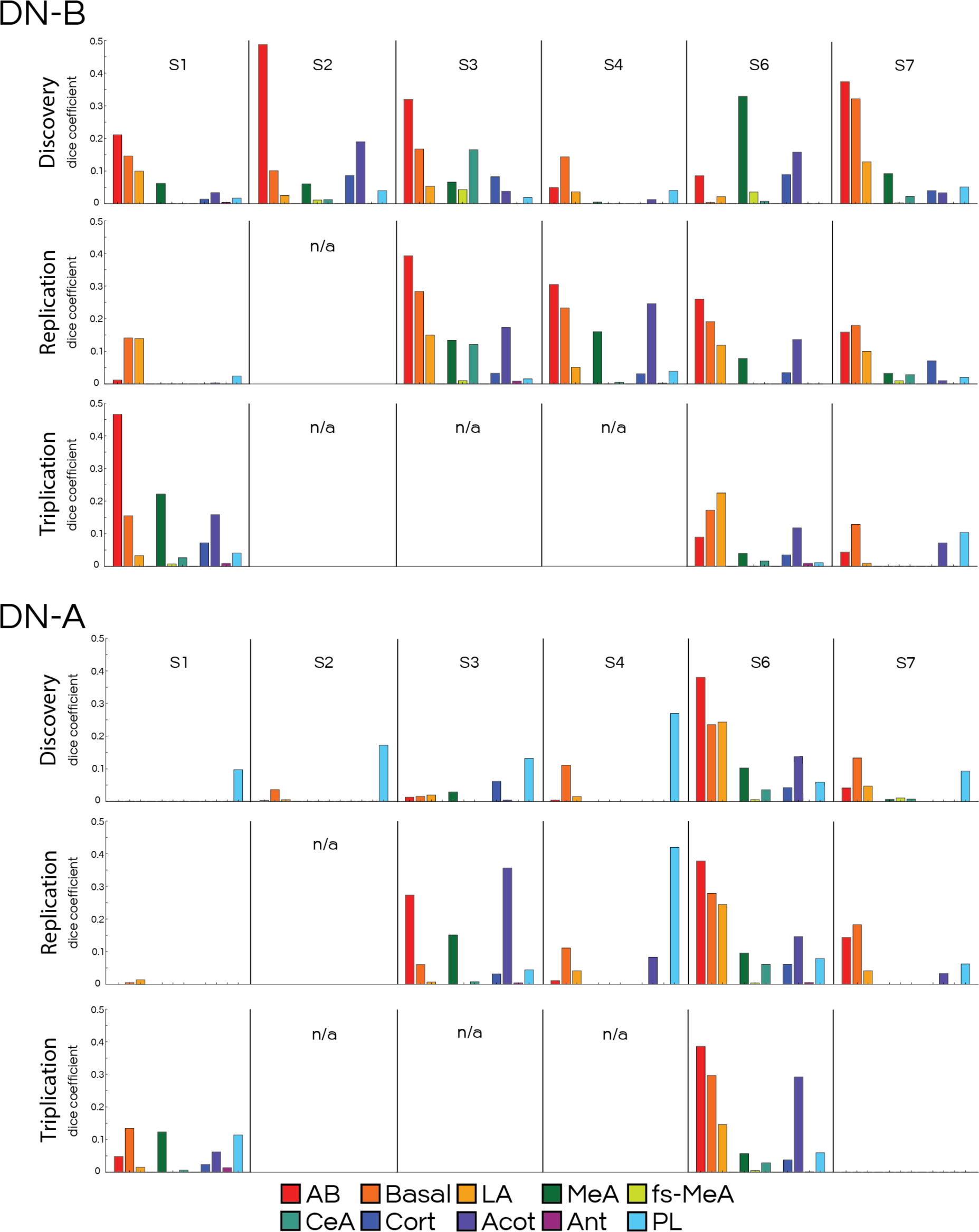
Quantification of the overlap between regions of default network B (DN-B) and an individualized segmentation of amygdala nuclei. In each subject, we used the Dice coefficient to calculate the overlap between the thresholded DN-B correlations (shown in Figs. 4 & 5) and the map of each amygdala nucleus defined using an automated, individualized segmentation procedure. We also assessed overlap with a hand-drawn map of the medial nucleus (MeA; shown in Fig. 6). In all individuals except subject S6, the DN-B regions overlapped mostly with the basolateral complex of the amygdala (red-yellow colors), which includes accessory basal (AB), basal, and lateral (LA) nuclei. This finding was replicated and triplicated in individuals that provided independent left-out data. In most cases, the DN-B regions overlapped predominantly with the accessory basal nucleus, followed by the nearby basal nucleus. When we repeated this analysis using the regions of DN-A (lower graphs), we did not observe the same pattern consistently, with the exception of subject S6. The remaining abbreviations refer to central (CeA), cortical (Cort), corticoamygdaloid (Acot), anterior (Ant), and paralaminar (PL) nuclei. Fs-MeA refers to the estimate of the medial nucleus that was created by FreeSurfer (as opposed to the hand-drawn estimate; MeA). “n/a” indicates participants who did not provide replication or triplication datasets.

**Supplementary Figure S5:**
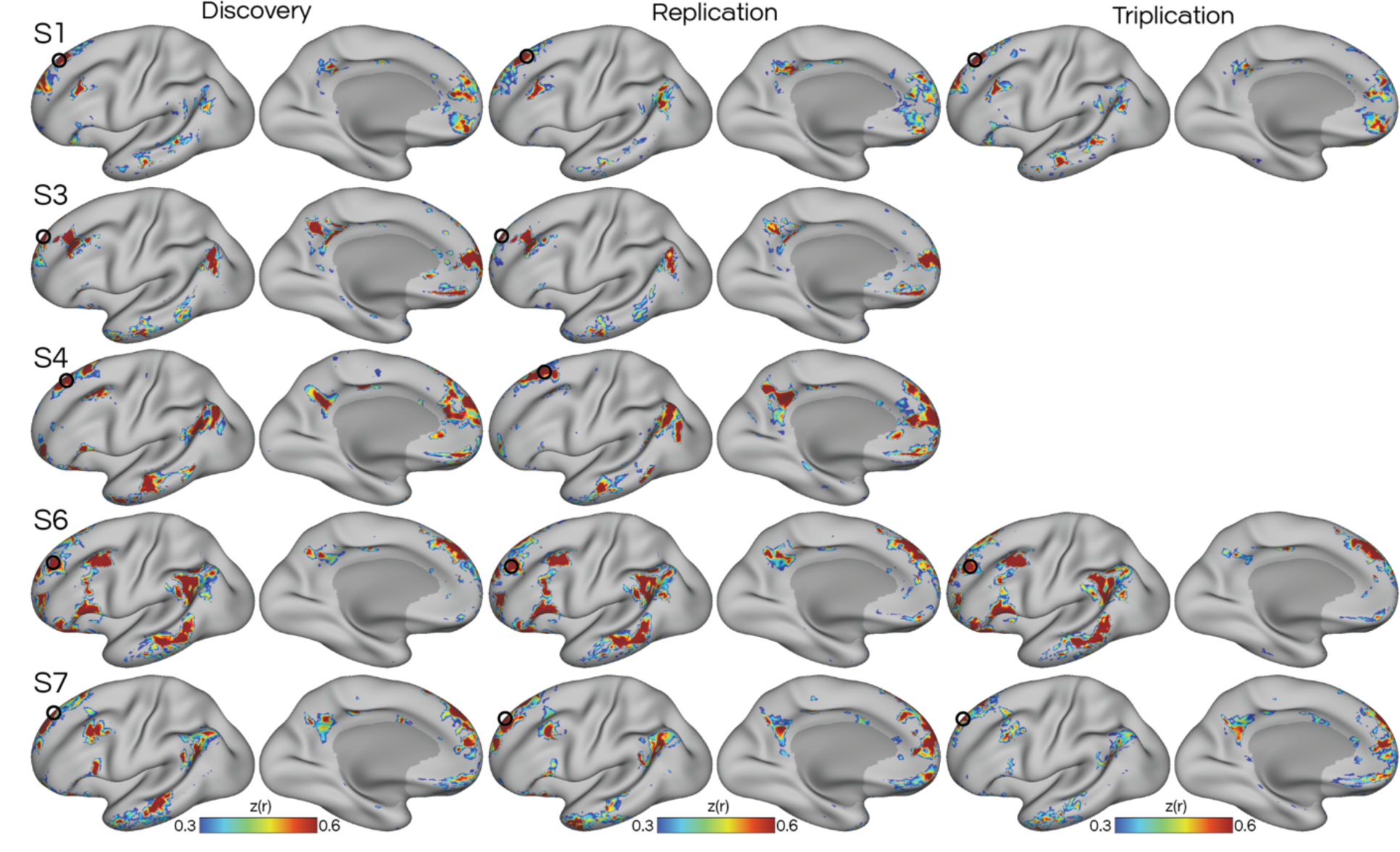
Replication and triplication of volume-defined default network B (DN-B) in left-out datasets. Once the analyses of the discovery dataset had been completed, definition of DN-B was replicated and triplicated in individuals who provided sufficient data. Networks were defined in the volume through manually selected seed voxels and then projected to the surface for visualization (similar to Fig. 1). Dorsolateral prefrontal seeds were selected in each dataset to target DN-B. In each dataset, DN-B occupied a similar distribution of regions and displayed similar separation from DN-A (not shown).

**Supplementary Figure S6:**
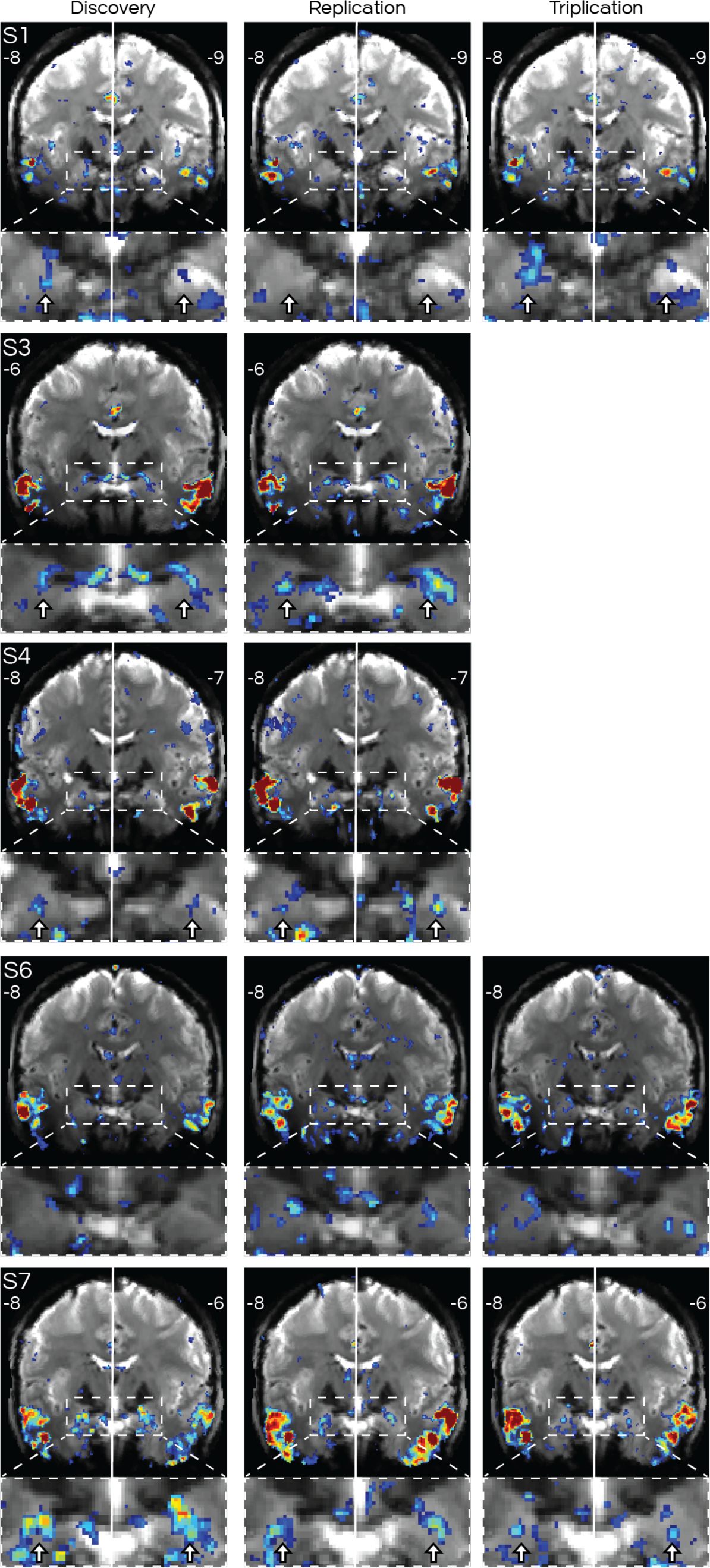
Replication and triplication of basolateral amygdala region of default network B (DN-B). Bilateral amygdala regions were reproducibly observed in 4 of the 5 subjects who provided held-out data, providing evidence that the amygdala contains circumscribed regions that show connections to network DN-B. Subject S2 is not included as they only provided a discovery dataset. Subject S1 showed limited evidence at this threshold in the replication dataset, but displayed clear regions in the triplication dataset. Subject S4 showed stronger evidence (correlations) in their replication dataset. For each subject, coronal slices are the same as those shown in Fig. 4. The dashed box indicates the location of the zoom-in underneath each full coronal slice. The white arrows indicate putative basolateral amygdala regions of DN-B that did not overlap with DN-A, and are consistent within subjects across panels, to serve as landmarks.

**Supplementary Figure S7:**
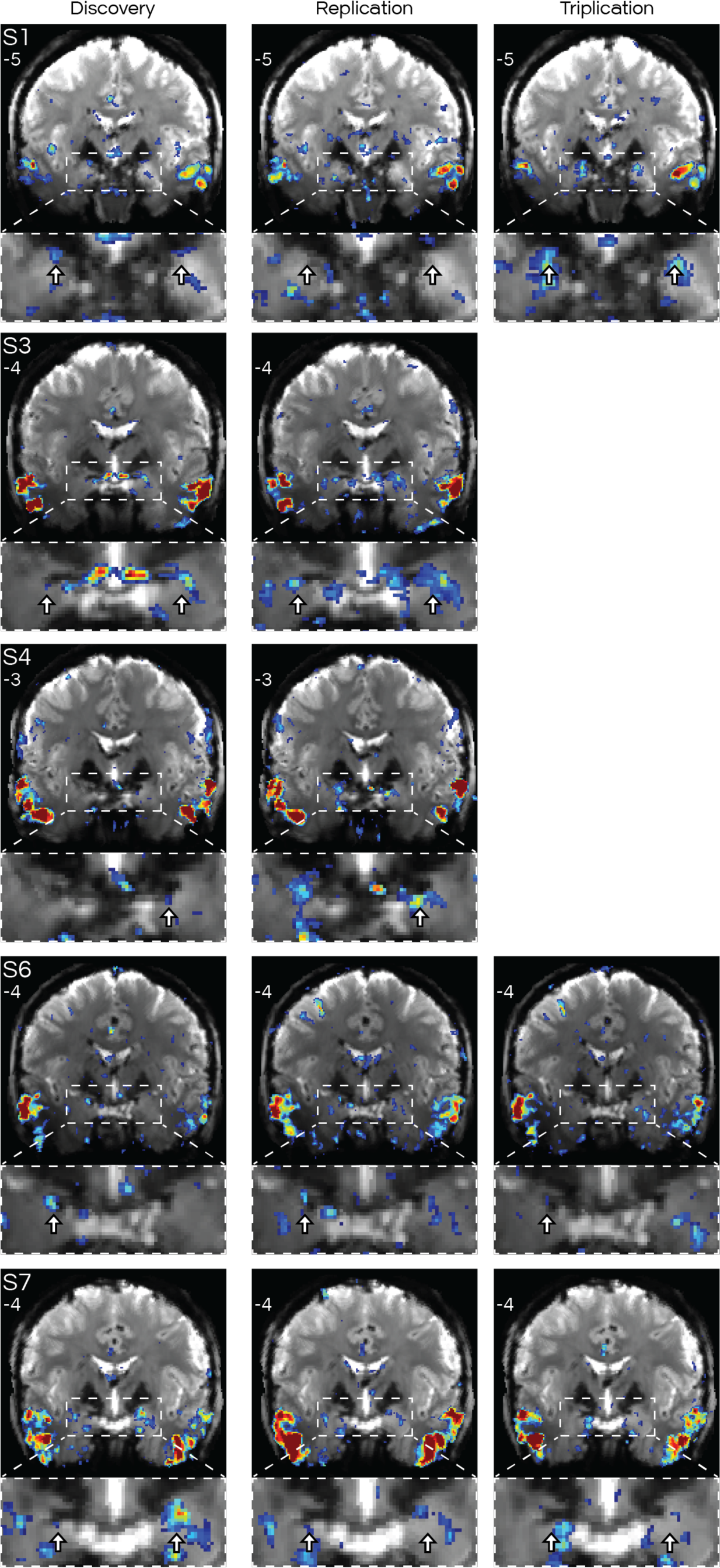
Replication and triplication of medial amygdala (MeA) regions of default network B (DN-B). Formatted according to Supp. Fig. S6. The presence of medial amygdala regions was replicated and triplicated in all individuals that provided left-out data. Two subjects (S4, S6) that appeared to only have unilateral MeA regions in their discovery dataset showed evidence of bilateral regions in replication and/or triplication. For each subject, coronal slices are the same as those shown in Fig. 4. White arrows indicate location of putative MeA regions of DN-B that did not overlap with DN-A, and are consisted within participants across panels to serve as landmarks.

**Supplementary Figure S8:**
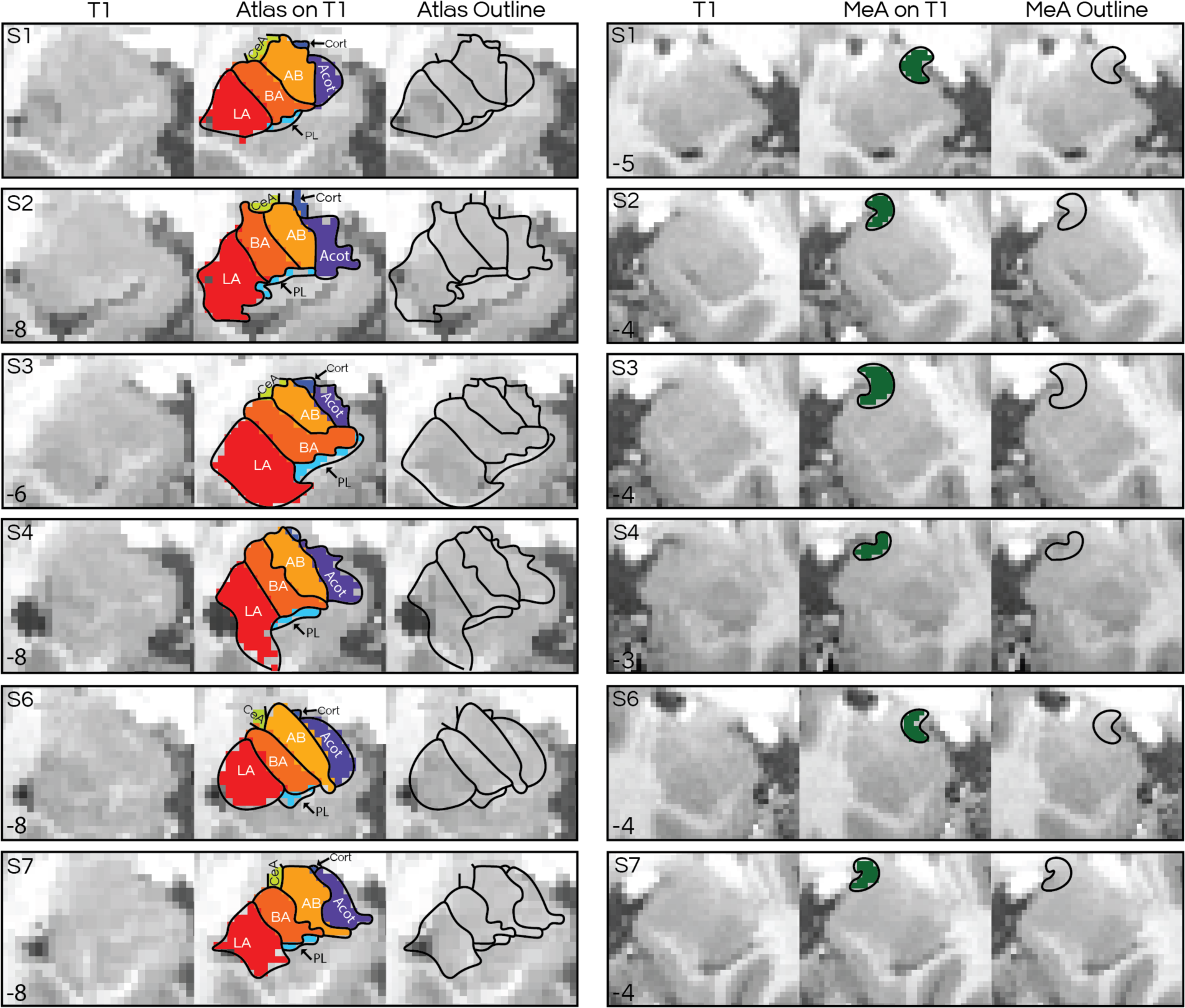
Images displaying the correspondence between anatomical (T1) images in each individual and individualized estimates of the amygdala nuclei. Coronal slices show the T1 anatomical image alongside individualized demarcations of the amygdala nuclei (colors) which were hand delineated (black lines) to serve as landmarks. Left columns show the automated, individualized segmentation obtained from FreeSurfer (“Atlas”), while the right panels show the hand-drawn estimate of the medial nucleus (“MeA”) drawn by a trained expert (author Q.Y.) who was blinded to the network maps. The estimates of the nuclei should be seen as approximations, rather than strict boundaries, given the difficulties in identifying demarcations between nuclei in neuroimaging data. Abbreviations are listed in Supp. Fig. S4.

